# Conservation strategy insights for three protected *Phengaris* butterflies combining genetic and landscape analyses

**DOI:** 10.1101/2025.03.12.642762

**Authors:** Jérémy Gauthier, Annabelle Sueur, Camille Pitteloud, Rémi Clément, Julia Bilat, Nadir Alvarez, Nicolas Gorius, Delphine Danancher

## Abstract

Biodiversity loss is accelerating globally, with insects among the most affected taxa. Wetlands, crucial ecosystems providing essential services, have experienced significant decline due to agricultural intensification and habitat destruction. These changes threaten species that rely on wetland habitats, such as the *Phengaris* butterflies, which exhibit complex ecological dependencies on specific host plants and ant species. Their populations are structured into metapopulations, where connectivity between habitat patches plays a crucial role in species persistence. This study focuses on population genomics and landscape analysis of three *Phengaris* species (*P. alcon, P. nausithous, P. teleius*) in the Bugey Mountain Massif, France, with the aim of developing a conservation strategy. Using ddRAD sequencing, we analyzed genetic diversity, population structure, and gene flow across multiple localities, revealing distinct population clusters with varying degrees of connectivity. Despite some populations exhibiting high genetic diversity levels, others demonstrate low heterozygosity and signs of genetic isolation, emphasizing conservation concerns. Migration estimates and resistance mapping further identified key dispersal corridors and isolated patches requiring targeted interventions. Our findings support metapopulation conservation strategies that prioritize maintaining connectivity, enhancing degraded patches, and identifying new potential habitats. By combining genomic and landscape data, we propose an evidence-based approach to conservation planning, facilitating adaptive management for *Phengaris* species. This study provides a practical framework for habitat restoration and corridor management in fragmented ecosystems, ensuring long-term species resilience in rapidly changing landscapes.

## Introduction

Global biodiversity is experiencing an unprecedented decline, with increasing evidence that insects are among the most affected taxa (Dirzo et al., 2014; Hallmann et al., 2017; Wagner et al., 2021). This decline is driven by multiple factors, with habitat destruction and agricultural intensification being among the primary causes ((Dirzo et al., 2014; Hallmann et al., 2017; Wagner et al., 2021)). Wetlands have been particularly affected throughout the 20th century (Fluet-Chouinard et al., 2023). Agricultural intensification has led to wetlands drying and conversion to agricultural land. Since the early 20th century, over 70% of European wetlands have been lost (Davidson, 2014), with an additional 45% disappearing since the 1970s (Hu et al., 2017). Those that remain are deeply impacted by anthropogenic pressures such as climate change, drainage, water level regulation, afforestation, and eutrophication (Brinson & Malvárez, 2002; Moomaw et al., 2018; Verhoeven, 2014). Due to their ecological vulnerability, wetlands are now protected under the Ramsar Convention and conservation initiatives at national and European levels, such as the Nature Restoration Act and the Habitats Directive.

Wetlands are biodiversity hotspots, providing approximately 40% of global ecosystem services (Okruszko et al., 2011). They host unique communities and the loss of these habitats has a direct impact on species that are strictly associated with them. This is the case for butterfly species of the *Phengaris* genus (Lepidoptera, Lycaenidae). In Europe, this genus, previously named *Maculinea*, encompasses three protected species: *Phengaris alcon* (Denis & Schiffermüller, 1775), *Phengaris nausithous* (Bergsträsser, 1779), and *Phengaris teleius* (Bergsträsser, 1779). These species are characterized by intricate ecological dependencies. Their life cycles need specific host plants and ant species, displaying a remarkable example of obligate myrmecophily (Als et al., 2004). Butterflies rely on specific environmental conditions, including meadows rich in diverse flora (Romo et al., 2015). The reliance of *Phengaris* species on specific plants and ant species complicates their conservation, necessitating targeted efforts to preserve both their habitats and associated biodiversity (Maes et al., 2024).

Due to their extreme specialization, *Phengaris* species exhibit patchy distributions and fluctuating population dynamics in metapopulations, as revealed in various studies (Boe et al., 2022; Jansen et al., 2012; Maes et al., 2024; Mouquet et al., 2005; Nowicki et al., 2019; Popović et al., 2017; Vanden Broeck et al., 2017). Metapopulations consist of local populations connected by occasional dispersal, where habitat patches may experience independent extinctions and recolonizations (as defined by (Hanski & Gilpin, 1991)). The balance of these processes determines species persistence. Thus, the size and genetic composition of groups colonizing new habitats, as well as dispersal rates between populations, play an essential role in metapopulation dynamics (Austin et al., 2011). In addition, habitat disturbance significantly influences these dynamics (Hanski et al., 1995, 2017). Firstly, the direct destruction of habitats will lead to local extinction of populations (DiLeo et al., 2024)). In addition, loss of genetic diversity at the metapopulation level, due to recurrent local extinctions and/or recolonization by related individuals, can increase its vulnerability to collapse globally. Secondly, the disruption of the network that exists between local populations will impact the metapopulation dynamics (Hanski et al., 1995). Studies highlighted that *Phengaris* species exhibit limited dispersal, strongly influenced by landscape structure (Nowicki et al., 2014; Popović et al., 2017; Skórka, Nowicki, Kudłek, et al., 2013; Skórka, Nowicki, Lenda, et al., 2013).

Genetic data has become an invaluable tool in conservation biology (Schmidt et al., 2023; Sucháčková Bartoňová et al., 2023; Willi et al., 2022), especially for species whose dynamics need to be finely characterized. Genetic data provides insights into population viability and sustainability. For example, low genetic diversity often indicates a reduced effective population size (either reflected by the current census size or, if not, suggesting a recent bottleneck) and a narrower capacity to adapt to disturbances (Hoffmann et al., 2017; L. F. Keller & Waller, 2002). Small and/or isolated populations are more subject to mechanisms that erode their genetic diversity through inbreeding and genetic drift ((Hoffmann et al., 2017; L. F. Keller & Waller, 2002)). Assessing population size and genetic diversity among different populations can help identify populations in decline, and thus prioritize conservation efforts. In the case of metapopulation dynamics, the study of population structure and gene flow is essential for designing protected areas and corridors that maintain or enhance genetic exchange (Sherpa et al., 2022). Genetic information allows a fine diagnosis of threatened populations, guiding conservation strategies that foster the long-term persistence and adaptability of species (Allendorf et al., 2010; Schmidt et al., 2023; Suchá čková Bartoň ová et al., 2023; Willi et al., 2022).

However, assessing populations genetic diversity and their connectivity requires accounting for landscape constraints (D. Keller et al., 2015; Segelbacher et al., 2010). This is the aim of landscape genetics, which combines landscape and genetic analyses to finely characterize the underlying constraints (Lowe & Allendorf, 2010; Manel et al., 2003). In recent years, new innovative approaches have been developed, in particular the integration of graph theory (Savary et al., 2024). These approaches focus on connectivity, conceptualizing it as a network composed of interconnected nodes (habitat patches) and links (dispersal paths) (Savary et al., 2022). These methods offer a complete and useful framework to support conservation strategies.

The context of the present study was the implementation of a conservation strategy for the three above-mentioned *Phengaris* species in the Bugey mountain massif in eastern France. The aim was to elaborate a scalable conservation strategy, identifying the habitat patches to be conserved in priority, as well as the connections to be restored or reinforced. To achieve this, we used an approach combining population genomics and landscape analysis. Using extensive sampling across occupied habitat patches, we applied ddRAD sequencing to generate population genomic data. In parallel, a detailed landscape analysis was performed to identify possible links between the various patches. This approach also defined likely dispersal areas in which new habitat patches could be promoted. Finally, the combination of these approaches enabled the creation of connectivity maps that reveal particularly useful for managers in developing a concrete, local conservation strategy for the three studied species.

## Material and methods

### Study area and monitoring

The study localities were determined based on known distribution of the species and their host plants, namely *Sanguisorba officinalis* for *P. teleius* and *P. nausithous* and *Gentiana pneumonanthe* for *P. alcon* (available data from the “Conservatoire Botanique National Alpin”, the “Syndicat du Haut-Rhône” and the “Conservatoire d’Espaces Naturels Rhône-Alpes” as part of the “Plan Ouest Lémanique pour la Connaissance et la Conservation des Azurés POLCCA”). A monitoring program of *Phengaris* populations initiated in 2017 is still ongoing. Given the small surface area of the study sites and the reduced population sizes, the monitoring method selected was a sight count based on the time spent prospecting. The monitoring sessions were carried out on sunny days with little or no wind and at least three times a year during the flight period of each species, in order to obtain a count as close as possible to the peak of emergence. Temporary captures were made to avoid double counting. Based on count data for the five years from 2017 to 2021, temporation effective population sizes (*N*e-t) were estimated by calculating the harmonic mean (Vucetich et al., 1997) (Supplementary Table 1).

### Genetic sampling

Genetic sampling has been performed during the summer 2020. In order to minimize the impact on individuals and populations, sampling was conducted as late as possible in the season, i.e. after the breeding season. In each locality, 12 samples were targeted (Table 1). For each individual, a leg was carefully removed and placed in absolute alcohol. The individual was then quickly released to minimize impact on the survival of captured individuals (Koscinski et al., 2011). Capture and sampling permit was delivered by the Direction régionale de l’environnement, de l’aménagement et du logement (DREAL) Auvergne-Rhône-Alpes (DDPP01-20-197). For each individual captured, the corresponding GPS position was recorded.

**Table 1.** Descriptive statistics on the genetic status estimated at both locality and population level. For each species, the number of samples and the number of SNPs are indicated, followed by the estimated values for: observed heterozygosity (Ho), mean gene diversities within population (Hs), inbreeding coefficient (*F*IS) and effective population size (Ne-g). Note that for small sampling sizes (<12 samples) the values are less reliable or could not be estimated.

| Species | Locality | N | Nb SNP | Ho (se) | Hs (se) | $F_{IS}$ (se) | $N_e$-g [CI] |
| --- | --- | --- | --- | --- | --- | --- | --- |
| *P. alcon* | Belloire | 12 | 7651 | 0.082 (0.002) | 0.093 (0.002) | 0.067 (0.006) | 70.3 [64.4 - 77.3] |
|  | Bidonnes | 4 |  | 0.109 (0.002) | 0.120 (0.002) | 0.038 (0.009) | NA |
|  | Cerin | 12 |  | 0.078 (0.002) | 0.088 (0.002) | 0.058 (0.005) | 21.8 [21.1 - 22.5] |
|  | Ormes | 12 |  | 0.046 (0.001) | 0.051 (0.001) | 0.037 (0.007) | 63 [55.2 - 73.3] |
| *P. nausithous* | Bidonnes | 12 | 11107 | 0.125 (0.002) | 0.133 (0.002) | 0.036 (0.004) | 34 [33.4 - 34.7] |
|  | Broues | 12 |  | 0.140 (0.002) | 0.150 (0.002) | 0.054 (0.004) | 44.4 [43.4 - 45.3] |
|  | Flon | 2 |  | 0.121 (0.003) | 0.157 (0.003) | 0.081 (0.012) | NA |
|  | merged | 26 |  | 0.131 (0.002) | 0.157 (0.002) | 0.111 (0.003) | 12.7 [12.6 - 12.7] |
|  | Epierre | 11 |  | 0.077 (0.001) | 0.085 (0.001) | 0.048 (0.004) | 9.4 [9.2 - 9.5] |
|  | Intriat | 12 |  | 0.063 (0.001) | 0.070 (0.001) | 0.050 (0.005) | 67.9 [63.2 - 73.3] |
|  | merged | 23 |  | 0.069 (0.001) | 0.087 (0.001) | 0.089 (0.004) | 5.3 [5.3 - 5.4] |
| *P. teleius* | Bidonnes | 20 | 11538 | 0.106 (0.002) | 0.119 (0.002) | 0.070 (0.004) | 51.4 [50.6 - 52.3] |
|  | Broues | 12 |  | 0.102 (0.002) | 0.117 (0.002) | 0.085 (0.005) | 29.7 [29.1 - 30.3] |
|  | Flon | 4 |  | 0.093 (0.002) | 0.092 (0.002) | -0.044 (0.009) | NA |
|  | merged | 36 |  | 0.103 (0.001) | 0.123 (0.001) | 0.098 (0.003) | 26 [25.9 - 26.1] |
|  | Lavours | 5 |  | 0.084 (0.002) | 0.092 (0.002) | 0.059 (0.008) | NA |
|  | Melogne | 4 |  | 0.047 (0.001) | 0.056 (0.001) | 0.075 (0.012) | NA |
|  | Vaux | 3 |  | 0.077 (0.002) | 0.091 (0.002) | 0.063 (0.010) | NA |
|  | merged | 12 |  | 0.070 (0.001) | 0.097 (0.001) | 0.189 (0.006) | 7.6 [7.5 - 7.7] |
|  | Epierre | 13 |  | 0.050 (0.001) | 0.051 (0.001) | 0.007 (0.004) | 46.2 [43.1 - 49.8] |
|  | Pont Loup | 12 |  | 0.051 (0.001) | 0.056 (0.001) | 0.056 (0.006) | 32.4 [31 - 34] |
|  | merged | 25 |  | 0.050 (0.001) | 0.069 (0.001) | 0.115 (0.004) | 1.5 [1.5 - 1.5] |

### ddRADseq

Genomic DNA was extracted from each leg using QIAamp DNA Micro kits (Qiagen) following the manufacturer’s instructions including a lysis step with proteinase K overnight at 56°C. DNA concentration was evaluated using a Qubit assay (Thermo Fisher Scientific). ddRAD libraries were prepared by digesting DNA with the restriction enzymes PstI and MseI. Individual-specific combinations of adapters and indexes were then linked to identify each sample. The libraries were amplified by PCR for 20 cycles, purified, quantified using a Picogreen assay (Thermo Fisher Scientific), and pooled equimolarly. Size selection with PippinPrep (SageScience) was performed to retain only fragments between 325 and 385 bp. Final libraries were sequenced using an Illumina HiSeq2500 125 bp paired-end at the Lausanne Genomic Technology Facility (LGTF).

### Populations genomics

Raw reads were demultiplexed with individual barcodes and cleaned to remove low-quality bases (qual<30) and technical sequences using CutAdapt 2.0 (Martin, 2011). Read quality was assessed using the FASTQC tool (Babraham Institute). For each species, ipyrad 0.9.84 (Eaton & Overcast, 2020) was then used with a clustering threshold of 90% to reconstruct loci and identify genetic variations. Genetic variations were filtered using vcftools 0.1.16 (Danecek et al., 2011) to retain only informative SNPs, *i*.*e*. bi-allelic SNPs shared by at least 80% of samples from each species.

A subset of independent SNPs was extracted to perform genetic structure inference, keeping only one SNP per locus. To investigate genetic population structure, two different approaches have been used. Population admixture was assessed using STRUCTURE 2.3.3 (Pritchard et al., 2000) and testing K-value from 1 to 10. For each value, three runs were performed with a burn-in of 100,000 followed by 500,000 iterations. The most likely number of clusters was determined using the Evanno method (Evanno et al., 2005) implemented in Structure Harvester (Earl & vonHoldt, 2012). As a complementary approach, principal component analysis (PCA) implemented for genetic data in the adegenet R package (Jombart & Ahmed, 2011) was used.

Two complementary approaches were applied to assess connectivity among localities. First, genetic differentiation (FST) was estimated between each locality pair using the hierfstat R package (Goudet, 2005). Second, relative migration rates were estimated using divMigrate from the diveRsity 1.9.90 package (Keenan et al., 2013) and 100 bootstrap iterations to calculate mean and 95% confidence intervals. Three indices have been estimated, Jost’s D (Jost, 2008), Nei’s Gst (Nei & Chesser, 1983), and the effective number of migrants Nm (Alcala et al., 2014).

To assess the genetic status of localities and populations, observed heterozygosities (Ho), observed gene diversities (He), allelic richness (Ar) and inbreeding coefficient (FIS) were estimated using the hierfstat R package (Goudet, 2005). To estimate population sizes using linkage disequilibrium information, loci were repositioned on the closest available reference genome which is the genome of *P. arion* (GCA_963565745.1) with the blastn tool (Camacho et al., 2009). Each SNP was then repositioned by considering the first base of the blast hit and the loci. An additional SNP filter was applied to exclude lowest allele frequency at 0.05 as rare alleles tend to bias estimations (Waples & Do, 2010). Genetic effective population sizes (*N*e-g) were estimated using LDNe (Waples & Do, 2008) at both locality and population scales. The two effective population size estimations, *i*.*e. N*e-g and *N*e-t, were then compared.

### Landscape analyses

To model the potential movement of individuals between different localities, we relied on graph theory and the methods implemented in the Graphab tool (Foltête et al., 2021). The land cover of the study areas was obtained by combining data from Grand Genève, the Syndicat de la rivière d’Ain aval et ses affluents (SR3A), and the Ain departmental inventory of eco-landscape continuities of interest (TVB01). Land uses were homogenized: first, their resolution, set at 5mx5m, but also certain occupations such as forests, meadows and edges, depending on whether or not they belong to wetlands. A buffer was applied to transport networks, *i*.*e*. roads and railroads, of 2m or 5m depending on their size (main or secondary network). Areas of overlap between the rasters were used to integrate habitat continuity. The resistance of these different habitats was assessed by combining information from the literature (www.trameverteetbleue.fr) and expert knowledge. Three types of habitat were identified: attractive habitats (species habitats and similar), elements that are preferentially skirted during movements (hedges, edges, etc.) and blocking elements (urbanized areas, large forests, large-scale farming). Friction coefficients ranging from 1 to 1000 were then assigned for each type of land cover (Supplementary Table 2). Based on these maps, dispersion simulations were performed using graph theory with the Graphab tool (Foltête et al., 2021). For each species, patches where their presence is attested were defined as nodes. Links correspond to the optimal path between patches and are calculated by optimizing the cumulative costs assigned to the pixels of the resistance matrix.

### Crossing of genetic and landscape graphs

To confirm or reject a path between two localities, the genetic structure of the populations was used. “Confirmed” paths connect two localities belonging to the same genetic cluster. Indeed, the observed population homogeneity indicates gene flow and individuals migrating among localities. Conversely, theoretical links between localities belonging to different genetic clusters were not considered. The maximum dispersal distance was set at the distance of the longest confirmed link for each species. Link fitting into the dispersal range but linking localities with no genetic information were considered “putative”. In addition, based on the dispersal distances, potential dispersal zones were estimated around each butterfly site, taking into account the resistance matrix.

## Results

### Genetic structure and gene flow

The ddRAD population genomics approach proved highly effective to identify the spatial genetic structure of the metapopulation, allowing the integration of all samples. Collecting a single leg provided sufficient DNA for population genomic analyses (Supplementary Table 3), while minimizing the impact on the populations studied. After filtering, we obtained 7,651 shared SNPs for *P. alcon*, 11,107 for *P. nausithous* and 11,538 for *P. teleius*, enabling us to perform in-depth genetic analyses.

A comparative genetic structure analysis revealed similarities between the three *Phengaris* species. For *P. alcon*, the four sampled localities form distinct genetic clusters, or populations in a metapopulation framework, as indicated by genetic structure analyses (PCA, admixture) and *F*ST estimates, with the lowest differentiation (*F*ST = 0.536) observed between Belloire and Cerin. The Ormes and Belloire localities appear geographically close by direct distance (Figure 2a), but show a strong genetic differentiation of *F*ST = 0.581. For *P. nausithous*, the five localities sampled are genetically structured into two populations. The first population is located at the northeast of the study area, while the second, comprising the Intriat and Epierre localities, is more centrally located (Figure 2D). Given the distribution of the differentiation values, we consider that the level of genetic differentiation is low between localities belonging to the same population with an *F*ST < 0.226, while it is higher between localities of different populations characterized by an *F*ST > 0.492 (Supplementary Table 4). For *P. teleius*, in addition to the northeastern and central-western populations, a third population appears in the southeast of the study area. It is not identified as a cluster by the STRUCTURE approach with the best K value of 2, but clearly forms a distinct population in the PCA, more specifically on the PC2 which carries 7.9% of the genetic variance explained. Estimates of differentiation show moderate values of *F*ST, but the low numbers of localities in this population suggest that these values should be considered with caution. The geographical distribution of the populations highlighted similarities among the three species, with one population in the northeast, close to Lake Geneva, and one in the center-west.

**Figure 1.**
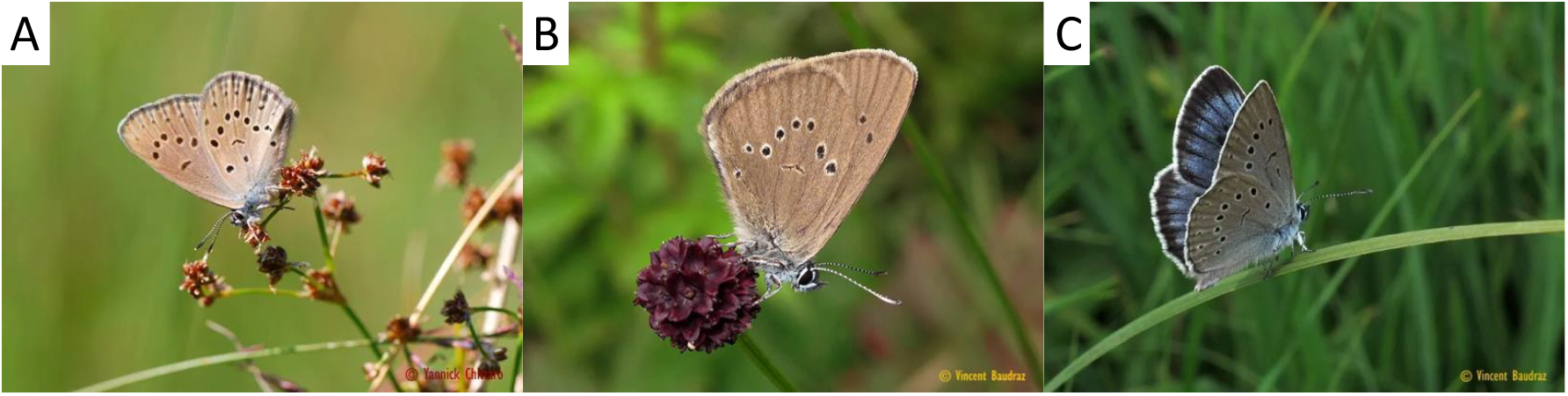
Habitus of the three studied species, A *P. alcon*, B *P. nausithous* and C *P. teleius* (credits Michel Baudraz)

**Figure 2.**
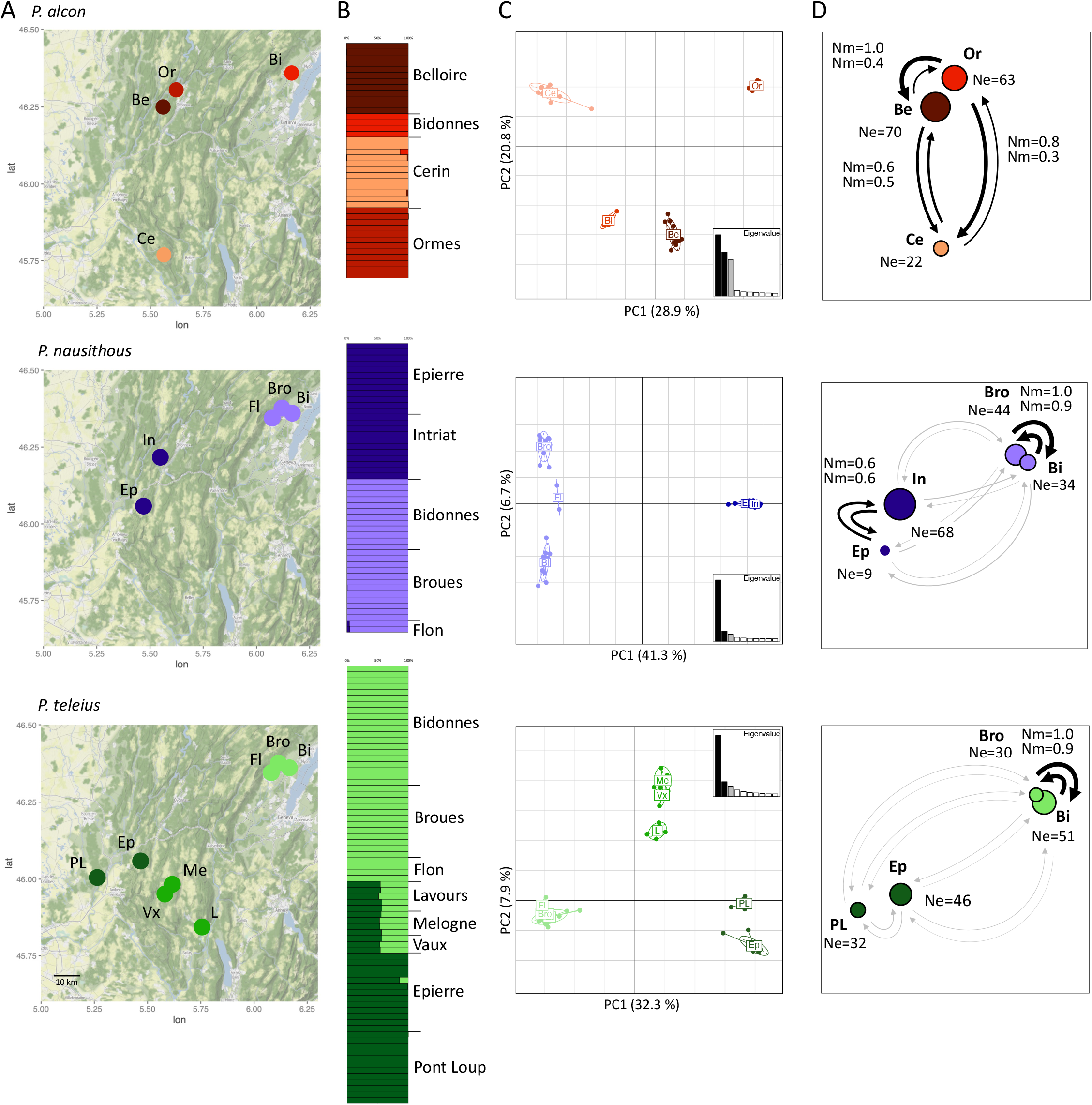
For each species, from top to down *P. alcon, P. nausithous* and *P. teleius*, distribution maps of sampled localities (A), genetic structure plots from STRUCTURE (B), PCA plots on genetic data (C), and schematic representation of the population descriptive statistics (D), *i*.*e*. effective population size (*N*e) and effective number of migrants (*N*m) for each locality with a sampling size supé rieur à 10. Dot size is proportional to sampling size (detailed in Supplementary Table 2). (Be = Belloire, Bi = Bidonnes, Bro = Broues, Ce = Cerin, Ep = Epierre, Fl = Flon, In = Intriat, L = Lavours, Me = Melogne, Or = Ormes, PL = Pont Loup, Vx = Vaux).

For *P. nausithous* and *P. teleius*, the results of migration levels between different localities are fully consistent with the observed genetic structure and estimated levels of genetic differentiation (Figure 2; Supplementary Table 4). The effective number of migrants (*N*m), which combines information from the other two statistics, Gst and D, and is the most appropriate for our dataset, shows a clear inverse correlation between migration levels and genetic differentiation. Logically, we observe a high level of migration between localities belonging to the same population, *N*m > 0.574, and a low level between localities of different populations, *N*m < 0.125 (Figure 2 D). It should be noted that a low level of migration was estimated between the Epierre and Pont-Loup localities, even though both localities belong to the same population, possibly revealing a former connectivity between the localities that has since been lost. Unexpectedly, *P. alcon* localities, despite being genetically structured, exhibit relatively high migration levels (*N*m > 0.326), even between the most geographically distant sites, such as Ormes and Cerin (Figure 2 D).

### Genetic diversity and population sizes

For each species, the localities to the north-east of the study area - Bidonnes, Broue and Belloire - show a significantly higher level of observed heterozygosity than the other localities. There is great variability in the heterozygosity observed among localities belonging to different populations, with differences ranging from simple to double. This is the case for Ormes and Belloire for *P. alcon*, Broue and Intriat for *P. nausithous* and Epierre and Bidonnes for *P. teleius* (Table 1). For each species, the localities at the north-east of the study area - Bidonnes, Broue and Belloire - show a significantly higher level of observed heterozygosity than the other localities. Population-level estimates for *P. nausithous* and *P. teleius, i*.*e*. by grouping samples from different localities according to the observed genetic structure, are consistent with locality-level estimates, and the observed values appear to correspond to the averages of the values estimated at the locality level.

The inbreeding coefficient (FIS) varies among localities. While a general inverse relationship with heterozygosity is observed, exceptions exist. For example, for *P. teleius*, the Epierre and Pont Loup localities, which belong to the same genetic population and have a similar level of heterozygosity, show a marked difference of the inbreeding coefficient. This may be linked to the larger effective population size of Epierre (Table 1). Population-level *F*IS estimates, obtained by merging samples from multiple localities, are consistently higher than locality-level estimates. This may be linked to the Wahlund effect and a population-scale substructure that represents the localities, which will tend to increase the inbreeding estimate (De Meeûs, 2018).

Genetic effective population sizes (*N*e-g) were estimated using linkage disequilibrium information, and only localities with a population size greater than 10 were included. For the three species, there was considerable heterogeneity in effective sizes between localities (Table 1). For *P. nausithous*, for example, there is a large difference between the two localities Epierre (*N*e-g = 9.4) and Intriat (*N*e-g = 67.9), which belong to the same genetic population. Estimates made at population level consistently reveal smaller effective sizes than estimates at locality level (Table 1), certainly linked to mixture-LD effect (Cox et al., 2024). It makes more sense to add up the local values, or to multiply the average of the local *N*e by the number of localities (Cox et al., 2024).

Comparison of effective sizes estimated on the basis of temporal counts (*N*e-t) in the previous five years using the harmonic mean reveals no correlation with genetic effective sizes (*N*e-g) based on linkage disequilibrium analysis for the three species (Pearson’s correlation test p-value > 0.05) (Figure 3A). The estimated size based on genetic information seems to be almost always slightly higher. Similarly, there is no correlation between genetic effective size (*N*e-g) and genetic diversity estimated from observed heterozygosity (Ho) (Pearson’s correlation test p-value > 0.05) (Figure 3B).

**Figure 3.**
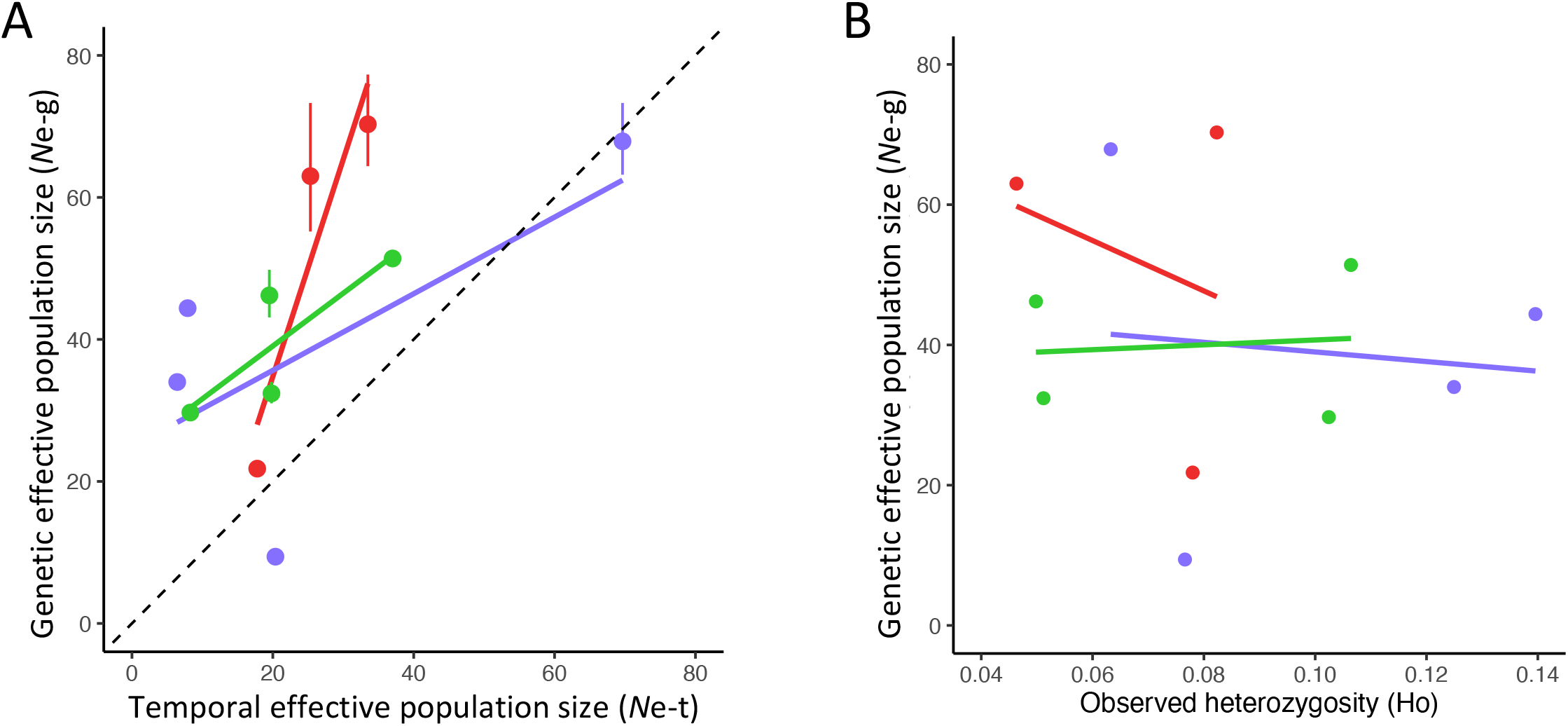
**A** Plot of the effective population sizes estimated using temporal counts (*N*e-t) and genetic information, *i*.*e*. linkage disequilibrium (*N*e-g). B Plot of the effective population size estimated using linkage disequilibrium and observed heterozygosity (Ho). *P. alcon* is represented in red, *P. nausithous* in purple and *P. teleius* in green.

### Landscape analyses and dispersal abilities

The resistance maps were used to estimate least-cost paths connecting pairs of patches with butterflies including patches for which we have genetic data. Using genetic data, we estimated the maximum dispersal distance as the greatest linear distance between two patches within the same genetic population, excluding cost-distance metrics due to their complexity in interpretation. These maximum dispersal distances were set at 18.5 km for *P. nausithous* and 18 km for *P. teleius*. For *P. alcon*, no maximum dispersal distance was determined, as all four genetically analyzed patches represent independent populations (Figure 2). In the integrative step, the use of the genetic information allowed us to correct the theoretical networks. Among putative dispersal paths, those exceeding the maximum dispersal distance were excluded. The other paths were of two types: “confirmed” paths, which connect patches belonging to the same genetic populations, and “putative” links, which also connect patches with butterflies for which we have no genetic information.

Network topology varies significantly across the study area (Figure 4, Supplementary Figure 1). In the northeast region (Figure 4 A,B), where *P. nausithous* and *P. teleius* are present in several patches, the networks are fairly extensive. *P. nausithous* showed a complex network with a high degree of redundancy and intersecting paths between patches. For *P. teleius* and in the same area, the absence of individuals observed in the south-eastern patch (in yellow on Figure 4 B) directly alters the density of the network. In contrast, the western network, which includes Pont Loup, Epierre and Intriat, is very elongated, with patches far apart (Figure 4 C,D). This is mainly due to Epierre, which is isolated in a more mountainous area. This locality connects with Intriat (*P. nausithous*) to the north and Pont Loup (*P. teleius*) to the west (Figure 4 C,D), but remains isolated from the southeastern network despite its proximity to certain localities (Supplementary Figure 1). However, the topology and the isolated setting of the Intriat locality seem to prevent gene flow with these localities.

**Figure 4.**
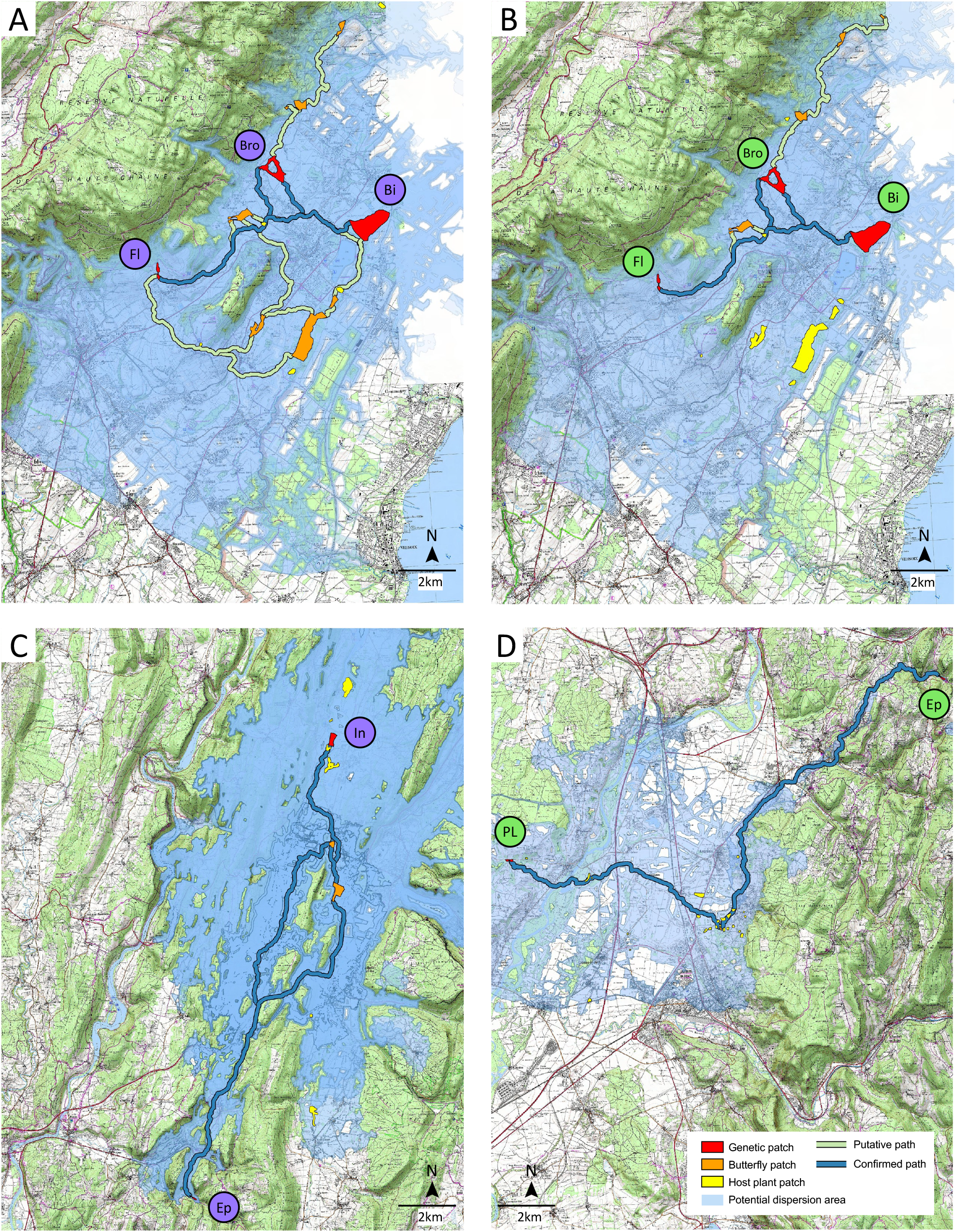
Decision map for *P. nausithous* (A, C) and *P. teleius* (B, C) including the paths identified between the genetic patches (in red) and the patches where the species is present (in orange). The maps also include the patches where the host plant is present (yellow) and the dispersal buffer around each genetic patch (blue).

Combining resistance maps and maximum dispersal distances, buffers were created around each species and genetic site to identify accessible areas on a landscape scale (shown in blue in Figure 4). These buffers can then be used to identify new patches of interest, *i*.*e*. host plant patches that can be found in the overlapping buffers of several butterfly matches. This is the case of the southern patches of the *P. teleius* network (Figure 4 B). By combining proximity to a network and the accessibility of host plant patches, it is possible to identify the most interesting and relevant patches to be favored in *Phengaris* conservation strategies.

## Discussion

### Current population state

The identification of a large number of molecular markers, over two thousands for each species, enabled the application of population genomics approaches to finely characterize the genetic state of populations as well as the connectivity among localities. The use of various complementary indices such as observed heterozygosity (Ho), inbreeding coefficient (FIS), migration rate (*N*m) and effective size (*N*e) reveals population dynamics. The highest observed heterozygosity values fall within the range of values observed locally in studies of butterflies in Europe (Després et al., 2019; Kebaïli et al., 2023; Trense et al., 2021) and even of species considered as endangered (Dupuis et al., 2020). However, some localities show diversities up to half as high, such as the Ormes locality for *P. alcon* or Epierres for *P. teleius*, reaching values close to those of almost extinct species (Nakahama et al., 2024). For conservation purposes, this information is crucial in selecting localities to be protected or supported by conservation or enhancement actions. Inbreeding levels, at both the locality and population scale, were lower than in the studies cited above. This represents a positive point for the conservation of the species. It could be linked to population dynamics and the strong gene flow observed between localities (Hanski et al., 1994).

The geographical distribution of genetic diversity showed shared patterns among the three species, with lower diversity levels in the western localities and greater diversity in the north-eastern localities. However, despite these northeastern localities, which include Bidonnes, Broues and Flon, are very isolated, they lie on the frontier between France and Switzerland and could therefore belong to a larger population encompassing additional localities from Switzerland not sampled in this study, mainly because of the difficulty of transferring genetic material of protected species across frontiers. Thus, comprehensive conservation strategies must integrate all these localities—both within France and in neighboring Switzerland—to effectively maintain genetic diversity and connectivity across the entire transboundary population..

The effective population sizes estimated at the local scale are quite small, with values ranging from 1.5 to 26. These values are largely below IUCN thresholds, such as the 50/500 rule (Jamieson & Allendorf, 2012), which are used to classify populations at high risk of extinction in the short-term (reviewed in Harmon & Braude, 2010). These findings reinforce concerns about these species and support their current protection status under Appendix II of the Bern Convention and Appendices II and IV of the Council of Europe’s Habitats Directive (1992).

### Meta-population dynamics

Genetic differentiation and migration rate analyses reveal strong gene flow between localities within the same genetic populations. This induces homogeneous genetic clusters as observed in the genetic structure analysis, but also seems to lead to average genetic diversity within populations. This strong gene flow plays a crucial role in counterbalancing the strong effect of local genetic drift, which is more intense in populations with very small numbers (Wang et al., 2016), as observed here. Given the small population sizes in each locality, insufficient gene flow would likely lead to rapid differentiation due to strong genetic drift. The strong intra-population gene flow is sometimes unexpected, as it concerns localities that are quite distant geographically. It is difficult to compare the cost-distances obtained from our modeling analyses with those from other studies, as the scale of friction coefficients differs greatly from one study to another. Thus, the maximum distances between two localities linked genetically are 18.5 km for *P. nausithous* and 18.0 km for *P. teleius*. These values are significantly greater than the dispersal distances known from the literature, which are always below 5km for the three species (Maes et al., 2004; Nowicki et al., 2005, 2014; Popović et al., 2017; Popović & Nowicki, 2023; Vanden Broeck et al., 2017). However, dispersal distances are generally estimated by mark-recapture approaches, which provide point-in-time data limited to the study area (Franzé n et al., 2024).

The results revealed meta-population dynamics as defined by (Hanski et al., 2017). Strong gene flow between localities from the same population tends to maintain and homogenize genetic diversity. The lack of correlation between genetic diversity and effective population sizes, already observed in the literature (Ryman et al., 2019) is the result of variability in local population sizes but homogenization of gene pools by gene flow. Population sizes vary both spatially, among localities, and temporally, as shown by the counting performed during the previous five years. The dynamics of the metapopulation involve a turnover from one year to the next and from one locality to another, depending on more or less favorable conditions (DiLeo et al., 2024). This metapopulation functioning has implications for conservation, which need to focus on protecting and strengthening the localities, but above all on maintaining the connectivity among them. This is the guarantee of a functional metapopulation that is characterized by the constant presence of favorable conditions and is thus able to maintain itself in the long term (Kral-O’Brien & Harmon, 2021; Sielezniew et al., 2019).

### Practical maps for conservation strategy

Genetic data provide detailed insights into population status, and when integrated with spatial analyses, can inform conservation strategies. For this reason, and in collaboration with the site managers, three strategies have been envisaged based on the results of the study.

#### Strengthening vulnerable patches

Genetic diversity, inbreeding, and effective population size analyses allow us to rank patches and identify those most at risk. The first strategy is therefore to try to increase conservation measures in these patches, either by directly increasing their surface area and habitat quality, or by identifying favorable patches in their close surroundings. The analysis of dispersal areas around the patches revealed potential new reachable patches. For example, this is the case for the Intriat locality occupied by *P. nausithous* (Figure 5C). This habitat patch has a small surface and the genetic diversity is the lowest of all those estimated for the species. However, surrounding habitat patches contain the host plant (in yellow on Figure 5C). Improving these habitats could allow the establishment of the species in these new patches even though presence of the host ant was not confirmed yet.

#### Strengthening the network by reinforcing existing paths

Regarding the connectivity among patches, detailed knowledge of the existing network is particularly informative in terms of the strategy to be adopted to maintain and strengthen this network. Genetic information has revealed connections between remote and isolated localities, e.g. Epierre and Pont Loup for *P. teleius* (Figure 5D), highlighting a degree of connectivity that earlier knowledge research did not predict. To maintain the existing network, connectivity can be enhanced by establishing new habitat patches along identified dispersal routes. More specifically regarding the later case, Pont Loup and Epierre are relatively distant, and the Epierre locality is isolated in the middle of a higher-altitude massif. However, analysis of the maps reveals that there is a cluster of favorable habitats between the two localities. Helping to establish a butterfly population halfway between these two localities would reinforce connectivity by creating a form of stepping-stone zone. It should be noted that in this process, only those localities for which a connection has been established by genetics should be supported. Facilitating gene flow between distinct populations should be avoided, as it may disrupt local adaptation by introducing maladaptive alleles (Derry et al., 2019).

#### Identification of alternative paths to improve network resilience to putative disruptions

The third strategy is to identify alternative routes to those highlighted. While habitat managers can modify specific patches, they cannot control the entire landscape. Anthropogenic activities can lead to changes in land use and the creation of barriers that prevent the dispersal of individuals from one patch to another. It therefore seems crucial to consider alternative paths with new intermediate stepping-stone patches to guarantee connectivity. Enhancing network complexity improves resilience to future changes. For example, this is the case of *P. teleius* in the northeast part of the study area (Figure 5B). There is also a large *P. teleius* patch at the south of the network (yellow in Figure 5B). The establishment of butterflies in this patch would make the network more complex and thus resilien, as is the case for *P. nausithous* in the same area (Figure 5A). We can thus see that the *P. nausithous* network is more prone to allowing gene flow on either side of the obstacle at the center of the network. This network is therefore more resilient, and connectivity will be maintained even if one of the paths is impacted by a future obstacle.

The findings of this study, particularly the decision maps, are actively being used by the Conservatory of Natural Spaces to prioritize conservation areas. Identified host plant patches are currently under assessment, with conservation measures being implemented. Through agreements with landowners, adaptive management strategies and new practices are being introduced to support butterfly populations. For instance, mowing schedules are adjusted to align with *Phengaris* biology, occurring either early in the season, before adult emergence, or later, after egg-laying and caterpillar descent into ant nests (Popović & Nowicki, 2023). Additionally, in key areas, sowing of the host plant *Sanguisorba officinalis* is planned to enhance habitat quality and ensure sufficient plant abundance. Overall, these applications highlight the practical value of integrating genetics with landscape analysis for conservation planning.

## Acknowledgments

We would particularly like to thank Yves Rozier and Cécile Guérin for the crucial sampling. We thank Cécile Racapé and Jessica Giraldi for updating the land use maps. We would like to thank the members of the steering committee who have closely followed the project since its inception: Flavia APE, Syndicat du Haut-Rhône, Syndicat de la rivière d’Ain aval et affluents, Entente interdépartementale Rhône-Alpes pour la démoustication, Pays de Gex Agglo. We thank the Agence de l’eau Rhône-Méditerranée-Corse, the Ain department and the Auvergne-Rhône-Alpes region for the funding and support they provided to this work. We thank the DREAL for the permit and the Alpine Ecology Laboratory (LECA) of Grenoble for the long-term storage of the DNA samples. We would also like to thank Paul Savary for his advice.

## Data Availability Statement

Raw genomic data can be found under NCBI BioProjects SUBXXXXX. The pipelines including custom scripts have been made available in the Github repository (XXXX).

## Figure captions

**Supplementary Figure 1**. Additional decision map for *P. teleius* including the Melogne, Lavours and Vaux localities. Paths identified between the genetic patches (in red) and the patches where the species is present (in orange) are represented. The maps also include the patches where the host plant is present (yellow) and the dispersal buffer around each genetic patch (blue).

**Supplementary Table 1**. Count data for each locality performed during the five years from 2017 to 2021, and temporation effective population sizes (*N*e-t).

**Supplementary Table 2**. Friction coefficients, from 1 to 1000, assigned to each type of land cover.

**Supplementary Table 3**. Descriptive statistics for each sample including the locality, the date of capture, the GPS position, the number of sequenced reads, the number of SNPs and the percentage of missing.

**Supplementary Table 4**. Pairwise differentiation level *F*ST and migration rates estimated between each locality of the three species, including the Jost’s D, Nei’s Gst, and the effective number of migrants *N*m. Values in bold indicate reliable estimates with sample sizes greater than 10.

