## Supplementary Material for "Conservation strategy insights for three protected *Phengaris* butterflies combining genetic and landscape analyses"

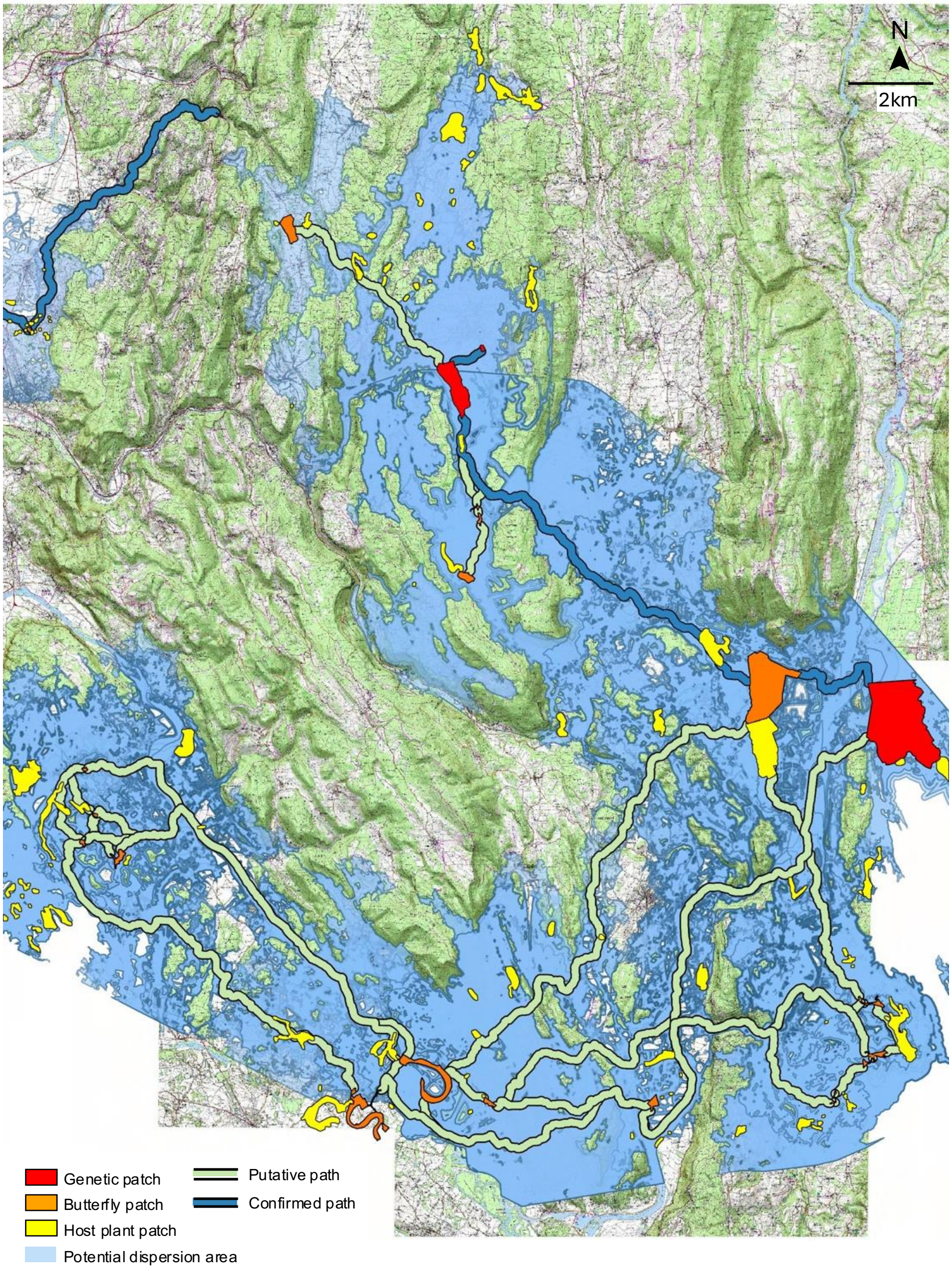


**Supplementary Figure 1.** Additional decision map for *P. teleius* including the Melogne, Lavours and Vaux localities. Paths identified between the genetic patches (in red) and the patches where the species is present (in orange) are represented. The maps also include the patches where the host plant is present (yellow) and the dispersal buffer around each genetic patch (blue).

**Supplementary Table 1.** Count data for each locality performed during the five years from 2017 to 2021, and temporation effective population sizes (*N*e-t).

| Species | Locality | 2017 | 2018 | 2019 | 2020 | 2021 | *N* e-t |
|---|---|---|---|---|---|---|---|
| *P. alcon* | Belloire | 120 | 90 | 30 | 15 | NA | 33.49 |
| | Bidonnes | 34 | 131 | 27 | 13 | 5 | 14.24 |
| | Cerin | 57 | 17 | 12 | 20 | 14 | 17.79 |
| | Ormes | 38 | NA | NA | 19 | NA | 25.33 |
| *P. nausithous* | Bidonnes | 4 | 3 | 10 | 28 | 17 | 6.43 |
| | Broues | 12 | 5 | 18 | 23 | 4 | 7.91 |
| | Flon | 4 | 11 | 13 | 2 | 2 | 3.53 |
| | merged | 20 | 19 | 41 | 53 | 23 | 26.40 |
| | Epierre | 81 | NA | 9 | NA | 42 | 20.37 |
| | Intriat | 46 | 52 | 122 | 121 | NA | 69.65 |
| | merged | 127 | 52 | 131 | 121 | 42 | 74.84 |
| *P. teleius* | Bidonnes | 26 | 51 | 40 | 46 | 33 | 37.01 |
| | Broues | 25 | 8 | 9 | 13 | 4 | 8.29 |
| | Flon | 25 | 29 | 7 | 5 | 2 | 5.45 |
| | merged | 76 | 88 | 56 | 64 | 39 | 59.78 |
| | Lavours | 41 | 14 | 23 | 26 | 199 | 27.35 |
| | Melogne | NA | 7 | 6 | 4 | NA | 5.36 |
| | Vaux | 46 | 17 | NA | 3 | 1 | 2.83 |
| | merged | 87 | 38 | 29 | 33 | 200 | 46.47 |
| | Epierre | 18 | NA | 28 | NA | 16 | 19.51 |
| | Pont Loup | NA | NA | 16 | 26 | NA | 19.81 |
| | merged | 18 | NA | 44 | 26 | 16 | 22.32 |

**Supplementary Table 2.** Friction coefficients, from 1 to 1000, assigned to each type of land cover.

| code | habitat name | value | 5455 | Edges of forest clearings | 10 | 742 | Spaces associated with communication networks | 100 |
|---|---|---|---|---|---|---|---|---|
| 1 | High-altitude meadows | 20 | 6000 | Trails and similar paths | 30 | 743 | Railway network | 100 |
| 2 | Dense urban area | 800 | 6219 | Water bodies over 100m wide | 300 | 811 | Industrial and commercial sites | 800 |
| 3 | Stream | 30 | 6317 | Intermittent watercourses with Burnet or Gentiana pneumonanthe | 1 | 811 | Sports and leisure infrastructure | 500 |
| 4 | Closed coniferous forests | 100 | 7119 | Continuous urban fabric over 100m wide | 1000 | 812 | Low-density buildings | 500 |
| 5 | Extensive meadows | 20 | 7119 | Monocultures over 10ha | 900 | 813 | Transport networks | 100 |
| 6 | Dry meadows | 30 | 8119 | Industrial and commercial sites over 100m wide | 1000 | 815 | Other impervious surfaces | 200 |
| 7 | Diffuse urban area | 500 | 8219 | Exclusive buildings in towns and villages over 100m wide | 1000 | 816 | Waste disposal site | 200 |
| 8 | Disturbed tree vegetation | 10 | 8227 | Permanent wet meadows with presence of Burnet or Gentians | 1 | 821 | Exclusive buildings in towns and villages | 800 |
| 9 | Closed deciduous forest | 100 | 8357 | Wetlands with Sanguisorba officinalis or Gentiana pneumonanthe | 1 | 821 | Permanent meadows | 20 |
| 10 | Intensive crops and meadows | 500 | 9129 | Temporary crops and meadows over 10ha | 900 | 822 | Abandoned urban and village buildings | 500 |
| 11 | Open forests | 30 | 9139 | Wet crops over 10ha | 900 | 822 | Permanent wet meadows | 2 |
| 12 | Other wetlands | 5 | 11100 | Open closed forests with pure conifers in patches | 30 | 823 | Dams | 800 |
| 13 | Calm waters | 30 | 11200 | Open closed forests with pure Scots pines | 30 | 825 | Vacant lots | 50 |
| 15 | Wet forests | 30 | 11259 | Closed forests with pure wet Scots pine over 100m wide | 500 | 831 | Meadows | 30 |
| 16 | Cliffs | 50 | 11300 | Open closed forests with fir or spruce | 30 | 831 | Organic waste | 500 |
| 17 | Wet meadows | 2 | 12100 | Open closed forests with predominant conifers and deciduous trees | 30 | 832 | Heathlands | 30 |
| 18 | River | 30 | 12159 | Closed forests with predominant conifers and wet deciduous trees over 100m wide | 500 | 833 | Scrublands | 30 |
| 19 | Vineyard | 30 | 21200 | Open closed forests with mixed deciduous trees | 30 | 835 | Wetlands | 5 |
| 20 | Orchard | 30 | 21259 | Closed forests with mixed wet deciduous trees over 100m wide | 500 | 911 | Vineyards and orchards | 30 |
| 21 | Railway | 100 | 21400 | Open closed forests with pure deciduous oaks | 30 | 912 | Temporary crops and meadows | 500 |
| 22 | Highway | 100 | 21459 | Closed forests with pure wet deciduous oaks over 100m wide | 500 | 913 | Wet crops | 500 |
| 23 | Road | 100 | 21500 | Open closed forests with pure deciduous trees in patches | 30 | 1010 | Intensive crops and meadows over 10ha | 900 |
| 24 | Gravel pit | 200 | 21559 | Closed forests with pure wet deciduous trees in patches over 100m wide | 500 | 1011 | Rocky environments | 50 |
| 25 | Paths | 30 | 21600 | Open closed forests with pure beech | 30 | 1111 | Heterogeneous agricultural areas | 20 |
| 28 | Urban vegetation | 500 | 21659 | Closed forests with pure wet beech over 100m wide | 500 | 1115 | Closed forests with pure wet conifers in patches | 100 |
| 49 | Closed coniferous forests over 100m wide | 1000 | 21700 | Open closed forests with pure black locust | 30 | 1119 | Closed forests with pure conifers in patches over 100m wide | 1000 |
| 99 | Closed deciduous forest over 100m wide | 1000 | 21800 | Open closed forests with pure other deciduous trees | 30 | 1119 | Still waters over 100m wide | 300 |
| 111 | Closed forests with pure conifers in patches | 100 | 21859 | Closed forests with pure wet other deciduous trees over 100m wide | 500 | 1125 | Closed forests with pure wet Scots pine | 100 |
| 111 | Still water | 30 | 22100 | Open closed forests with predominant deciduous trees and conifers | 30 | 1129 | Closed forests with pure Scots pine over 100m wide | 1000 |
| 112 | Closed forests with pure Scots pine | 100 | 22159 | Closed forests with predominant deciduous trees and wet conifers over 100m wide | 500 | 1139 | Closed forests with fir or spruce over 100m wide | 1000 |
| 112 | Floating vegetation | 20 | 23100 | Open closed forests without tree cover | 30 | 1215 | Closed forests with predominant conifers and wet deciduous trees | 30 |
| 113 | Closed forests with fir or spruce | 100 | 23159 | Closed forests without tree cover in wet areas over 100m wide | 100 | 1219 | Artificial surface waters over 100m wide | 400 |
| 113 | Oxbows | 30 | 31100 | Open forests with pure conifers | 30 | 1219 | Closed forests with predominant conifers and deciduous trees over 100m wide | 1000 |
| 114 | Flowing water | 20 | 32100 | Open forests with pure deciduous trees | 30 | 2117 | Raised bogs with presence of Gentians (P. alcon) or Burnet | 1 |
| 119 | Open forests over 100m wide | 500 | 32159 | Open forests with pure wet deciduous trees over 100m wide | 100 | 2125 | Closed forests with mixed wet deciduous trees | 30 |
| 121 | Artificial surface waters | 50 | 33100 | Open forests with mixed deciduous and conifers | 30 | 2127 | Low marsh with Gentians (P. alcon) or Burnet | 1 |
| 121 | Closed forests with predominant conifers and deciduous trees | 100 | 41200 | Open closed forests with mixed conifers | 30 | 2129 | Closed forests with mixed deciduous trees over 100m wide | 1000 |
| 127 | Other wetlands with Burnet or Gentians | 1 | 41400 | Open closed forests with pure Douglas fir | 30 | 2135 | Edges of reed beds | 10 |
| 131 | Aquatic vegetation | 20 | 41600 | Open closed forests with Corsican pine or pure black pine | 30 | 2137 | Reed beds with presence of Gentians (P. alcon) or Burnet | 1 |
| 139 | Calm waters over 100m wide | 300 | 41759 | Closed forests with wet fir or spruce over 100m wide | 500 | 2139 | Closed forests with pure chestnut over 100m wide | 1000 |
| 159 | Wet forests over 100m wide | 500 | 41800 | Open closed forests with pure conifers other than pine | 30 | 2145 | Closed forests with pure wet deciduous oaks | 30 |
| 211 | Raised bogs | 5 | 51105 | Edges of open wet forests | 10 | 2147 | Large sedge beds with presence of Gentians (P. alcon) or Burnet | 1 |
| 212 | Closed forests with mixed deciduous trees | 100 | 51109 | Open wet forest over 100m wide | 500 | 2149 | Closed forests with pure deciduous oaks over 100m wide | 1000 |
| 212 | Low marsh | 5 | 52105 | Edges of open deciduous forests | 10 | 2155 | Closed forests with pure wet deciduous trees in patches | 30 |
| 213 | Closed forests with pure chestnut | 100 | 52109 | Open deciduous forest over 100m wide | 1000 | 2159 | Closed forests with pure deciduous trees in patches over 100m wide | 1000 |
| 213 | Reed beds | 20 | 52205 | Open edges of birch and aspen grove | 10 | 2165 | Closed forests with pure wet beech | 30 |
| 214 | Closed forests with pure deciduous oaks | 100 | 52305 | Open edges of wet birch groves | 10 | 2175 | Closed forests with pure wet black locust | 30 |
| 214 | Large sedge beds | 5 | 52405 | Open edges of shrubland | 10 | 2179 | Closed forests with pure black locust over 100m wide | 1000 |
| 215 | Closed forests with pure deciduous trees in patches | 100 | 53105 | Edges of open coniferous forests | 10 | 2185 | Closed forests with pure wet other deciduous trees | 30 |
| 216 | Closed forests with pure beech | 100 | 53109 | Open coniferous forest over 100m wide | 1000 | 2189 | Closed forests with pure other deciduous trees over 100m wide | 1000 |
| 217 | Closed forests with pure black locust | 100 | 53205 | Open edges of pine forests | 10 | 2215 | Closed forests with predominant deciduous trees and wet conifers | 30 |
| 218 | Closed forests with pure other deciduous trees | 100 | 53209 | Open pine forest over 100m wide | 100 | 2217 | Wet meadow with presence of Gentians (P. alcon) or Burnet | 1 |
| 221 | Closed forests with predominant deciduous trees and conifers | 100 | 53305 | Open edges of peatland pine forests | 10 | 2219 | Closed forests with predominant deciduous trees and conifers over 100m wide | 1000 |
| 221 | Wet meadow | 2 | 54105 | Open edges of deciduous plantations | 10 | 2315 | Closed forests without tree cover in wet areas | 30 |
| 231 | Closed forests without tree cover | 50 | 54109 | Open deciduous plantation over 100m wide | 500 | 3119 | Open forests with pure conifers over 100m wide | 100 |
| 311 | Dry grasslands | 30 | 54205 | Edges of open poplar groves | 10 | 3215 | Open forests with pure wet deciduous trees | 30 |
| 311 | Open forests with pure conifers | 30 | 54305 | Open edges of wet poplar groves | 10 | 3219 | Open forests with pure deciduous trees over 100m wide | 100 |
| 321 | Mesic grasslands - meadows | 20 | 54405 | Open edges of coppices and small woods | 10 | 3319 | Open forests with mixed deciduous and conifers over 100m wide | 100 |
| 321 | Open forests with pure deciduous trees | 30 | 54505 | Open edges of forest clearings | 10 | 4110 | Open closed forests with mixed other conifer | 30 |
| 331 | Mesic meadows | 20 | 111500 | Open closed forests with pure wet conifers in patches | 30 | 4119 | Closed forests with mixed other conifers over 100m wide | 1000 |
| 331 | Open forests with mixed deciduous and conifers | 30 | 111900 | Open closed forests with pure conifers in patches over 100m wide | 1000 | 4125 | Closed forests with mixed wet conifers | 100 |
| 332 | Artificial grasslands | 50 | 112500 | Open closed forests with pure Scots pine in wet areas | 30 | 4129 | Closed forests with mixed conifers over 100m wide | 1000 |
| 341 | Open forests without tree cover | 50 | 112900 | Open closed forests with pure Scots pines over 100m wide | 1000 | 4139 | Closed forests with pure pine stands over 100m wide | 1000 |
| 341 | Parks and gardens | 50 | 113900 | Open closed forests with fir or spruce over 100m wide | 1000 | 4149 | Closed forests with pure Douglas fir over 100m wide | 1000 |
| 411 | Closed forests with mixed other conifers | 100 | 121100 | Open hedges | 10 | 4169 | Closed forests with pure Corsican pine or black pine over 100m wide | 1000 |
| 411 | Mesic thickets | 50 | 121500 | Open closed forests with predominant conifers and wet deciduous trees | 30 | 4175 | Closed forests with wet fir or spruce | 100 |
| 412 | Closed forests with mixed conifers | 100 | 121900 | Open closed forests with predominant conifers and deciduous trees over 100m wide | 1000 | 4179 | Closed forests with fir or spruce over 100m wide | 1000 |
| 413 | Closed forests with pure pine stands | 100 | 212500 | Open closed forests with mixed wet deciduous trees | 30 | 4189 | Closed forests with pure conifer other than pine over 100m wide | 1000 |
| 414 | Closed forests with pure Douglas fir | 100 | 212900 | Open closed forests with mixed deciduous trees over 100m wide | 1000 | 5000 | Forest edges | 10 |
| 416 | Closed forests with pure Corsican pine or black pine | 100 | 214500 | Open closed forests with pure wet deciduous oaks | 30 | 5005 | Wet forest edges | 10 |
| 418 | Closed forests with pure conifer other than pine | 100 | 214900 | Open closed forests with pure deciduous oaks over 100m wide | 1000 | 5110 | Open wet forest | 30 |
| 421 | Poplar groves | 30 | 215500 | Open closed forests with pure wet deciduous trees in patches | 30 | 5115 | Edge of wet forest | 10 |
| 421 | Wet heathlands (with M. caerulea) | 5 | 215900 | Open closed forests with pure deciduous trees in patches over 100m wide | 1000 | 5119 | Wet forest over 100m wide | 500 |
| 422 | Wet thickets | 30 | 216500 | Open closed forests with pure wet beeches | 30 | 5210 | Open deciduous forest | 30 |
| 431 | Hedges | 10 | 216900 | Open closed forests with pure beech over 100m wide | 1000 | 5215 | Edge of deciduous forest | 10 |
| 441 | Vines and fruit shrubs | 30 | 217500 | Open closed forests with pure wet black locust | 30 | 5219 | Deciduous forest over 100m wide | 1000 |
| 511 | Wet forest | 30 | 217900 | Open closed forests with pure black locust over 100m wide | 1000 | 5220 | Open birch and aspen grove | 30 |
| 521 | Deciduous forest | 100 | 218500 | Open closed forests with pure other wet deciduous trees | 30 | 5225 | Edge of birch and aspen grove | 10 |
| 522 | Birch and aspen grove | 30 | 218900 | Open closed forests with pure other deciduous trees over 100m wide | 1000 | 5229 | Birch and aspen grove over 100m wide | 100 |
| 523 | Wet birch groves | 10 | 221500 | Open closed forests with predominant deciduous trees and wet conifers | 30 | 5230 | Open wet birch groves | 30 |
| 524 | Shrubland | 30 | 221900 | Open closed forests with predominant deciduous trees and conifers over 100m wide | 1000 | 5235 | Edge of wet birch groves | 10 |
| 531 | Coniferous forests | 100 | 311900 | Open forests with pure conifers over 100m wide | 100 | 5240 | Open shrubland | 30 |
| 532 | Pine forests | 30 | 321500 | Open forests with pure wet deciduous trees | 30 | 5245 | Edge of shrubland | 10 |
| 533 | Peatland pine forests | 10 | 321900 | Open forests with pure deciduous trees over 100m wide | 100 | 5310 | Open coniferous forests | 30 |
| 541 | Deciduous plantations | 100 | 331900 | Open forests with mixed deciduous and conifers over 100m wide | 100 | 5315 | Edge of coniferous forest | 10 |
| 543 | Wet poplar groves | 10 | 412500 | Open closed forests with wet mixed conifers | 30 | 5319 | Coniferous forest over 100m wide | 1000 |
| 544 | Coppices and small woods | 100 | 412900 | Open closed forests with mixed conifers over 100m wide | 1000 | 5320 | Open pine forests | 30 |
| 545 | Forest clearings | 30 | 414900 | Open closed forests with pure Douglas fir over 100m wide | 1000 | 5325 | Edges of pine forests | 10 |
| 611 | Basins | 30 | 416900 | Open closed forests with Corsican pine or Douglas fir over 100m wide | 1000 | 5329 | Pine forest over 100m wide | 100 |
| 611 | Mineral surface | 50 | 417500 | Open closed forests with wet fir or spruce | 30 | 5330 | Open peatland pine forests | 10 |
| 621 | Inactive quarries | 200 | 417900 | Open closed forests with fir or spruce over 100m wide | 1000 | 5335 | Edges of peatland pine forests | 10 |
| 621 | Water bodies | 30 | 421500 | Open wet poplar groves | 30 | 5410 | Open deciduous plantations | 30 |
| 631 | Active quarries | 200 | 500000 | Edges of open forests | 10 | 5415 | Edges of deciduous plantations | 10 |
| 631 | Intermittent watercourses | 5 | 500500 | Edges of open wet forests | 10 | 5419 | Deciduous plantation over 100m wide | 500 |
| 632 | Permanent watercourses | 30 | 1215900 | Open closed forests with predominant conifers and wet deciduous trees over 100m wide | 500 | 5420 | Open poplar groves | 30 |
| 711 | Continuous urban fabric | 800 | 2125900 | Open closed forests with mixed wet deciduous trees over 100m wide | 500 | 5425 | Edges of poplar groves | 10 |
| 712 | Extensive monocultures | 200 | 2145900 | Open closed forests with pure wet deciduous oaks over 100m wide | 500 | 5430 | Open wet poplar groves | 10 |
| 721 | Market gardening | 200 | 2155900 | Open closed forests with pure wet deciduous trees in patches over 100m wide | 500 | 5435 | Edges of wet poplar groves | 10 |
| 721 | Material extraction sites | 200 | 2165900 | Open closed forests with pure wet beech over 100m wide | 500 | 5440 | Open coppices and small woods | 30 |
| 731 | Discontinuous urban fabric | 500 | 2185900 | Open closed forests with pure other wet deciduous trees over 100m wide | 500 | 5445 | Edges of coppices and small woods | 10 |
| 731 | Wastelands | 50 | 3215900 | Open forests with pure wet deciduous trees over 100m wide | 500 | 5449 | Coppices and small woods over 100m wide | 500 |
| 741 | Road network | 100 | 4175900 | Open closed forests with wet fir or spruce over 100m wide | 500 | 5450 | Open forest clearings | 30 |

**Supplementary Table 3.** Descriptive statistics for each sample including the locality, the date of capture, the GPS position, the number of sequenced reads, the number of SNPs and the percentage of missing.

| | | Geographic data | | | | Genetic data | | |
|---|---|---|---|---|---|---|---|---|
| Species | Code | Date | Long | Lat | Locality | Reads | SNP | %missing |
| *P. alcon* | Be-A01 | 30.07.2020 | 5.559081 | 46.2459 | Belloire | 364922 | 4656 | 18.80 |
| *P. alcon* | Be-A02 | 30.07.2020 | 5.559095 | 46.245776 | Belloire | 263902 | 3989 | 30.43 |
| *P. alcon* | Be-A03 | 30.07.2020 | 5.558988 | 46.245858 | Belloire | 342692 | 4236 | 26.12 |
| *P. alcon* | Be-A04 | 30.07.2020 | 5.557842 | 46.246514 | Belloire | 1219982 | 5612 | 2.13 |
| *P. alcon* | Be-A05 | 30.07.2020 | 5.559048 | 46.245936 | Belloire | 1973646 | 5496 | 4.15 |
| *P. alcon* | Be-A06 | 30.07.2020 | 5.558082 | 46.246319 | Belloire | 1457532 | 5619 | 2.01 |
| *P. alcon* | Be-A07 | 30.07.2020 | 5.558106 | 46.247014 | Belloire | 398312 | 4262 | 25.67 |
| *P. alcon* | Be-A08 | 30.07.2020 | 5.558242 | 46.246508 | Belloire | 1604600 | 5545 | 3.30 |
| *P. alcon* | Be-A09 | 14.08.2020 | 5.558205 | 46.246193 | Belloire | 950340 | 5063 | 11.70 |
| *P. alcon* | Be-A10 | 14.08.2020 | 5.558083 | 46.246361 | Belloire | 1440890 | 5692 | 0.73 |
| *P. alcon* | Be-A11 | 14.08.2020 | 5.557936 | 46.246399 | Belloire | 1101816 | 5225 | 8.88 |
| *P. alcon* | Be-A12 | 14.08.2020 | 5.558117 | 46.246335 | Belloire | 1229562 | 5654 | 1.40 |
| *P. alcon* | Bi-A01 | 21.07.2020 | 6.165564 | 46.364431 | Bidonnes | 1534036 | 5579 | 2.70 |
| *P. alcon* | Bi-A02 | 21.07.2020 | 6.1657 | 46.364597 | Bidonnes | 2397714 | 5373 | 6.30 |
| *P. alcon* | Bi-A03 | 21.07.2020 | 6.165164 | 46.36428 | Bidonnes | 1936138 | 5482 | 4.39 |
| *P. alcon* | Bi-A04 | 21.07.2020 | 6.164884 | 46.364511 | Bidonnes | 410106 | 4682 | 18.35 |
| *P. alcon* | Ce-A01 | 29.07.2020 | 5.565016 | 45.774696 | Cerin | 1696670 | 5637 | 1.69 |
| *P. alcon* | Ce-A02 | 29.07.2020 | 5.562839 | 45.775562 | Cerin | 1131856 | 5419 | 5.49 |
| *P. alcon* | Ce-A03 | 29.07.2020 | 5.562466 | 45.774926 | Cerin | 1646716 | 5196 | 9.38 |
| *P. alcon* | Ce-A04 | 29.07.2020 | 5.562854 | 45.775636 | Cerin | 1624530 | 5605 | 2.25 |
| *P. alcon* | Ce-A05 | 29.07.2020 | 5.563007 | 45.775934 | Cerin | 668392 | 5538 | 3.42 |
| *P. alcon* | Ce-A06 | 29.07.2020 | 5.563234 | 45.776495 | Cerin | 1580952 | 5665 | 1.20 |
| *P. alcon* | Ce-A07 | 29.07.2020 | 5.563023 | 45.775825 | Cerin | 234264 | 3294 | 42.55 |
| *P. alcon* | Ce-A08 | 29.07.2020 | 5.563118 | 45.775865 | Cerin | 1683414 | 5574 | 2.79 |
| *P. alcon* | Ce-A09 | 29.07.2020 | 5.562911 | 45.775526 | Cerin | 1394796 | 5599 | 2.35 |
| *P. alcon* | Ce-A10 | 29.07.2020 | 5.562777 | 45.777148 | Cerin | 1245166 | 5514 | 3.84 |
| *P. alcon* | Ce-A11 | 29.07.2020 | 5.562947 | 45.77528 | Cerin | 1919868 | 5623 | 1.94 |
| *P. alcon* | Ce-A12 | 29.07.2020 | 5.562934 | 45.775502 | Cerin | 1049154 | 5606 | 2.23 |
| *P. alcon* | Or-A01 | 14.08.2020 | 5.617659 | 46.307801 | Ormes | 1136400 | 5485 | 4.34 |
| *P. alcon* | Or-A02 | 14.08.2020 | 5.617207 | 46.307815 | Ormes | 1436672 | 5452 | 4.92 |
| *P. alcon* | Or-A03 | 14.08.2020 | 5.617494 | 46.307829 | Ormes | 953812 | 5576 | 2.76 |
| *P. alcon* | Or-A04 | 14.08.2020 | 5.61757 | 46.307864 | Ormes | 2341310 | 5650 | 1.46 |
| *P. alcon* | Or-A05 | 14.08.2020 | 5.61782 | 46.308018 | Ormes | 2087946 | 5426 | 5.37 |
| *P. alcon* | Or-A06 | 14.08.2020 | 5.616662 | 46.3081 | Ormes | 2067368 | 5567 | 2.91 |
| *P. alcon* | Or-A07 | 14.08.2020 | 5.616391 | 46.30806 | Ormes | 1001622 | 5280 | 7.92 |
| *P. alcon* | Or-A08 | 14.08.2020 | 5.61642 | 46.30808 | Ormes | 1711272 | 5710 | 0.42 |
| *P. alcon* | Or-A09 | 14.08.2020 | 5.616523 | 46.308033 | Ormes | 684960 | 5451 | 4.94 |
| *P. alcon* | Or-A10 | 14.08.2020 | 5.617591 | 46.307833 | Ormes | 1660734 | 5566 | 2.93 |
| *P. alcon* | Or-A11 | 14.08.2020 | 5.617505 | 46.307879 | Ormes | 1764806 | 5644 | 1.57 |
| *P. alcon* | Or-A12 | 14.08.2020 | 5.617918 | 46.307915 | Ormes | 502342 | 5243 | 8.56 |
| *P. nausithous* | Bi-N01 | 21.07.2020 | 6.165678 | 46.364627 | Bidonnes | 1289458 | 7540 | 7.71 |
| *P. nausithous* | Bi-N02 | 21.07.2020 | 6.165719 | 46.364542 | Bidonnes | 1614582 | 7924 | 3.01 |
| *P. nausithous* | Bi-N03 | 21.07.2020 | 6.165735 | 46.364512 | Bidonnes | 1994068 | 7857 | 3.83 |
| *P. nausithous* | Bi-N04 | 21.07.2020 | 6.165665 | 46.364722 | Bidonnes | 1677866 | 7977 | 2.36 |
| *P. nausithous* | Bi-N05 | 21.07.2020 | 6.165561 | 46.364496 | Bidonnes | 2160214 | 8046 | 1.52 |
| *P. nausithous* | Bi-N06 | 05.08.2020 | 6.165631 | 46.364567 | Bidonnes | 1974724 | 7902 | 3.28 |
| *P. nausithous* | Bi-N07 | 05.08.2020 | 6.165629 | 46.364488 | Bidonnes | 1542448 | 7802 | 4.50 |
| *P. nausithous* | Bi-N08 | 05.08.2020 | 6.165736 | 46.364545 | Bidonnes | 1515344 | 7563 | 7.43 |
| *P. nausithous* | Bi-N09 | 05.08.2020 | 6.165783 | 46.364518 | Bidonnes | 1796418 | 7870 | 3.67 |
| *P. nausithous* | Bi-N10 | 05.08.2020 | 6.165783 | 46.364484 | Bidonnes | 1272394 | 7749 | 5.15 |
| *P. nausithous* | Bi-N11 | 05.08.2020 | 6.165678 | 46.36462 | Bidonnes | 4234798 | 8019 | 1.85 |
| *P. nausithous* | Bi-N12 | 05.08.2020 | 6.165541 | 46.364617 | Bidonnes | 2486646 | 7998 | 2.11 |
| *P. nausithous* | Bro-N01 | 06.08.2020 | 6.129272 | 46.375364 | Broues | 1251858 | 7076 | 13.39 |
| *P. nausithous* | Bro-N02 | 06.08.2020 | 6.130762 | 46.374998 | Broues | 1891990 | 7516 | 8.00 |
| *P. nausithous* | Bro-N03 | 06.08.2020 | 6.131049 | 46.374668 | Broues | 1493894 | 7907 | 3.22 |
| *P. nausithous* | Bro-N04 | 06.08.2020 | 6.131464 | 46.374673 | Broues | 1332326 | 7152 | 12.46 |
| *P. nausithous* | Bro-N05 | 06.08.2020 | 6.131522 | 46.374644 | Broues | 1834732 | 7951 | 2.68 |
| *P. nausithous* | Bro-N06 | 06.08.2020 | 6.131633 | 46.374703 | Broues | 1158640 | 7168 | 12.26 |
| *P. nausithous* | Bro-N07 | 06.08.2020 | 6.131479 | 46.374786 | Broues | 2104026 | 7766 | 4.94 |
| *P. nausithous* | Bro-N08 | 06.08.2020 | 6.13142 | 46.374795 | Broues | 1928030 | 8022 | 1.81 |
| *P. nausithous* | Bro-N09 | 06.08.2020 | 6.131552 | 46.374752 | Broues | 2077490 | 7946 | 2.74 |
| *P. nausithous* | Bro-N10 | 06.08.2020 | 6.129853 | 46.375073 | Broues | 2197330 | 8024 | 1.79 |
| *P. nausithous* | Bro-N11 | 06.08.2020 | 6.129311 | 46.375365 | Broues | 2730830 | 7930 | 2.94 |
| *P. nausithous* | Bro-N12 | 06.08.2020 | 6.125176 | 46.37768 | Broues | 2648392 | 7996 | 2.13 |
| *P. nausithous* | Ep-N01 | 27.07.2020 | 5.467739 | 46.061752 | Epierre | 1578512 | 8086 | 1.03 |
| *P. nausithous* | Ep-N02 | 27.07.2020 | 5.467758 | 46.061793 | Epierre | 2188528 | 7820 | 4.28 |
| *P. nausithous* | Ep-N03 | 27.07.2020 | 5.467652 | 46.061786 | Epierre | 2196762 | 8049 | 1.48 |
| *P. nausithous* | Ep-N04 | 27.07.2020 | 5.466724 | 46.062377 | Epierre | 1946108 | 7843 | 4.00 |
| *P. nausithous* | Ep-N05 | 27.07.2020 | 5.46737 | 46.062037 | Epierre | 608930 | 6579 | 19.47 |
| *P. nausithous* | Ep-N06 | 27.07.2020 | 5.467595 | 46.061911 | Epierre | 460254 | 6330 | 22.52 |
| *P. nausithous* | Ep-N07 | 27.07.2020 | 5.467702 | 46.061836 | Epierre | 1642590 | 7406 | 9.35 |
| *P. nausithous* | Ep-N09 | 27.07.2020 | 5.46697 | 46.062242 | Epierre | 1766628 | 7611 | 6.84 |
| *P. nausithous* | Ep-N10 | 27.07.2020 | 5.467111 | 46.062152 | Epierre | 2319866 | 7932 | 2.91 |
| *P. nausithous* | Ep-N11 | 27.07.2020 | 5.467095 | 46.062174 | Epierre | 1241020 | 7669 | 6.13 |
| *P. nausithous* | Ep-N12 | 27.07.2020 | 5.467726 | 46.061788 | Epierre | 1456674 | 7279 | 10.91 |
| *P. nausithous* | Fl-N01 | 25.07.2020 | 6.086744 | 46.353214 | Flon | 1544192 | 7842 | 4.01 |
| *P. nausithous* | Fl-N02 | 25.07.2020 | 6.0863 | 46.354616 | Flon | 1909858 | 7893 | 3.39 |
| *P. nausithous* | In-N01 | 30.07.2020 | 5.545035 | 46.218976 | Intriat | 2251730 | 7409 | 9.31 |
| *P. nausithous* | In-N02 | 30.07.2020 | 5.545089 | 46.218991 | Intriat | 1154838 | 7717 | 5.54 |
| *P. nausithous* | In-N03 | 30.07.2020 | 5.545033 | 46.21931 | Intriat | 1988916 | 8086 | 1.03 |
| *P. nausithous* | In-N04 | 30.07.2020 | 5.545031 | 46.219323 | Intriat | 1636298 | 7830 | 4.16 |
| *P. nausithous* | In-N05 | 30.07.2020 | 5.544605 | 46.218869 | Intriat | 1248202 | 7813 | 4.37 |
| *P. nausithous* | In-N06 | 30.07.2020 | 5.544563 | 46.218843 | Intriat | 1300864 | 7654 | 6.32 |
| *P. nausithous* | In-N07 | 30.07.2020 | 5.544624 | 46.218835 | Intriat | 1217826 | 7553 | 7.55 |
| *P. nausithous* | In-N08 | 30.07.2020 | 5.544677 | 46.218725 | Intriat | 1079606 | 7973 | 2.41 |
| *P. nausithous* | In-N09 | 30.07.2020 | 5.544503 | 46.21883 | Intriat | 1811270 | 7950 | 2.69 |
| *P. nausithous* | In-N10 | 30.07.2020 | 5.54492 | 46.218574 | Intriat | 1600028 | 7880 | 3.55 |
| *P. nausithous* | In-N11 | 30.07.2020 | 5.544936 | 46.218569 | Intriat | 1812756 | 8012 | 1.93 |
| *P. nausithous* | In-N12 | 30.07.2020 | 5.544873 | 46.218692 | Intriat | 1208852 | 7822 | 4.26 |
| *P. teleius* | Bi-A05 | 21.07.2020 | 6.165372 | 46.3646 | Bidonnes | 233434 | 4415 | 46.90 |
| *P. teleius* | Bi-A06 | 05.08.2020 | 6.163463 | 46.364262 | Bidonnes | 774406 | 7317 | 11.99 |
| *P. teleius* | Bi-A07 | 05.08.2020 | 6.164448 | 46.364731 | Bidonnes | 1011952 | 7545 | 9.25 |
| *P. teleius* | Bi-A08 | 05.08.2020 | 6.165017 | 46.364926 | Bidonnes | 1398706 | 8108 | 2.48 |
| *P. teleius* | Bi-A09 | 05.08.2020 | 6.164071 | 46.364432 | Bidonnes | 993310 | 4855 | 41.60 |
| *P. teleius* | Bi-A10 | 05.08.2020 | 6.16561 | 46.36428 | Bidonnes | 2100602 | 8190 | 1.49 |
| *P. teleius* | Bi-A11 | 05.08.2020 | 6.16324 | 46.363779 | Bidonnes | 1544016 | 8051 | 3.16 |
| *P. teleius* | Bi-A12 | 05.08.2020 | 6.163614 | 46.364116 | Bidonnes | 1803058 | 8139 | 2.10 |
| *P. teleius* | Bi-T01 | 21.07.2020 | 6.163312 | 46.363596 | Bidonnes | 1600656 | 8202 | 1.35 |
| *P. teleius* | Bi-T02 | 21.07.2020 | 6.16407 | 46.363775 | Bidonnes | 2113654 | 7783 | 6.39 |
| *P. teleius* | Bi-T03 | 21.07.2020 | 6.163405 | 46.363924 | Bidonnes | 832998 | 7809 | 6.07 |
| *P. teleius* | Bi-T04 | 21.07.2020 | 6.162796 | 46.364077 | Bidonnes | 1766842 | 8199 | 1.38 |
| *P. teleius* | Bi-T05 | 21.07.2020 | 6.165006 | 46.363605 | Bidonnes | 1333956 | 7920 | 4.74 |
| *P. teleius* | Bi-T06 | 21.07.2020 | 6.165466 | 46.364615 | Bidonnes | 1621712 | 8149 | 1.98 |
| *P. teleius* | Bi-T07 | 21.07.2020 | 6.165272 | 46.36438 | Bidonnes | 456388 | 6427 | 22.70 |
| *P. teleius* | Bi-T07_x | 05.08.2020 | 6.165466 | 46.364615 | Bidonnes | 1184218 | 7928 | 4.64 |
| *P. teleius* | Bi-T08 | 21.07.2020 | 6.165289 | 46.364318 | Bidonnes | 1272302 | 7946 | 4.43 |
| *P. teleius* | Bi-T09 | 21.07.2020 | 6.163396 | 46.364242 | Bidonnes | 2006190 | 8063 | 3.02 |
| *P. teleius* | Bi-T10 | 21.07.2020 | 6.164669 | 46.364565 | Bidonnes | 3769928 | 8235 | 0.95 |
| *P. teleius* | Bi-T12 | 21.07.2020 | 6.165068 | 46.364344 | Bidonnes | 3679822 | 8225 | 1.07 |
| *P. teleius* | Bro-T01 | 06.08.2020 | 6.125643 | 46.377881 | Broues | 1456570 | 8175 | 1.67 |
| *P. teleius* | Bro-T02 | 06.08.2020 | 6.125219 | 46.377723 | Broues | 1863538 | 8081 | 2.80 |
| *P. teleius* | Bro-T03 | 06.08.2020 | 6.131456 | 46.374873 | Broues | 1577836 | 8080 | 2.81 |
| *P. teleius* | Bro-T04 | 06.08.2020 | 6.131429 | 46.374911 | Broues | 2059198 | 8092 | 2.67 |
| *P. teleius* | Bro-T05 | 06.08.2020 | 6.130009 | 46.374984 | Broues | 1613184 | 8081 | 2.80 |
| *P. teleius* | Bro-T06 | 06.08.2020 | 6.130292 | 46.374964 | Broues | 1849884 | 8223 | 1.09 |
| *P. teleius* | Bro-T07 | 06.08.2020 | 6.130064 | 46.374978 | Broues | 1301484 | 7888 | 5.12 |
| *P. teleius* | Bro-T08 | 06.08.2020 | 6.129276 | 46.375442 | Broues | 911624 | 7655 | 7.93 |
| *P. teleius* | Bro-T09 | 06.08.2020 | 6.129231 | 46.375428 | Broues | 2139348 | 8048 | 3.20 |
| *P. teleius* | Bro-T10 | 06.08.2020 | 6.125397 | 46.377856 | Broues | 609714 | 6663 | 19.86 |
| *P. teleius* | Bro-T11 | 06.08.2020 | 6.130858 | 46.374802 | Broues | 1241786 | 7986 | 3.95 |
| *P. teleius* | Bro-T12 | 06.08.2020 | 6.130811 | 46.374775 | Broues | 2086916 | 8076 | 2.86 |
| *P. teleius* | Ep-N08 | 27.07.2020 | 5.466931 | 46.062284 | Epierre | 2200896 | 8244 | 0.84 |
| *P. teleius* | Ep-T01 | 27.07.2020 | 5.467753 | 46.06174 | Epierre | 355944 | 5475 | 34.15 |
| *P. teleius* | Ep-T02 | 27.07.2020 | 5.467319 | 46.062129 | Epierre | 1254258 | 7251 | 12.79 |
| *P. teleius* | Ep-T03 | 27.07.2020 | 5.467892 | 46.061653 | Epierre | 1960112 | 7951 | 4.37 |
| *P. teleius* | Ep-T04 | 27.07.2020 | 5.467726 | 46.061816 | Epierre | 1676148 | 7456 | 10.32 |
| *P. teleius* | Ep-T05 | 27.07.2020 | 5.466804 | 46.06232 | Epierre | 1637656 | 8058 | 3.08 |
| *P. teleius* | Ep-T06 | 27.07.2020 | 5.467984 | 46.061559 | Epierre | 2301486 | 8200 | 1.37 |
| *P. teleius* | Ep-T07 | 12.08.2020 | 5.467216 | 46.062059 | Epierre | 1021240 | 7939 | 4.51 |
| *P. teleius* | Ep-T08 | 12.08.2020 | 5.467704 | 46.061794 | Epierre | 2242832 | 8075 | 2.87 |
| *P. teleius* | Ep-T09 | 12.08.2020 | 5.467742 | 46.061756 | Epierre | 1522878 | 8167 | 1.77 |
| *P. teleius* | Ep-T10 | 12.08.2020 | 5.467777 | 46.061671 | Epierre | 1657430 | 8213 | 1.21 |
| *P. teleius* | Ep-T11 | 12.08.2020 | 5.468256 | 46.061377 | Epierre | 1980494 | 8136 | 2.14 |
| *P. teleius* | Ep-T12 | 12.08.2020 | 5.468252 | 46.061438 | Epierre | 829090 | 7807 | 6.10 |
| *P. teleius* | Fl-T01 | 25.07.2020 | 6.086413 | 46.354262 | Flon | 1687168 | 8138 | 2.12 |
| *P. teleius* | Fl-T02 | 25.07.2020 | 6.086513 | 46.353526 | Flon | 1640388 | 8019 | 3.55 |
| *P. teleius* | Fl-T03 | 25.07.2020 | 6.086607 | 46.354118 | Flon | 1669450 | 8078 | 2.84 |
| *P. teleius* | Fl-T04 | 25.07.2020 | 6.086497 | 46.354124 | Flon | 1895356 | 7934 | 4.57 |
| *P. teleius* | L-T01_x | 10.08.2020 | 5.74402 | 45.836674 | Lavours | 1810412 | 7941 | 4.49 |
| *P. teleius* | L-T02_x | 10.08.2020 | 5.74402 | 45.836674 | Lavours | 1223046 | 7980 | 4.02 |
| *P. teleius* | L-T03_x | 10.08.2020 | 5.74402 | 45.836674 | Lavours | 1191576 | 8033 | 3.38 |
| *P. teleius* | L-T04_x | 10.08.2020 | 5.74402 | 45.836674 | Lavours | 729066 | 7838 | 5.73 |
| *P. teleius* | L-T05_x | 17.08.2020 | 5.74402 | 45.836674 | Lavours | 1515656 | 7978 | 4.04 |
| *P. teleius* | Me-T01 | 29.07.2020 | 5.606966 | 45.969083 | VauxMelogne | 2241582 | 7964 | 4.21 |
| *P. teleius* | Me-T02 | 29.07.2020 | 5.606938 | 45.969167 | VauxMelogne | 481660 | 7073 | 14.93 |
| *P. teleius* | Me-T03 | 29.07.2020 | 5.606851 | 45.969139 | VauxMelogne | 1749740 | 8127 | 2.25 |
| *P. teleius* | Me-T04 | 29.07.2020 | 5.606747 | 45.969074 | VauxMelogne | 794690 | 7359 | 11.49 |
| *P. teleius* | PL-T01 | 27.07.2020 | 5.261479 | 46.005952 | PontLoup | 1338522 | 8131 | 2.20 |
| *P. teleius* | PL-T02 | 27.07.2020 | 5.261214 | 46.005912 | PontLoup | 1186534 | 8146 | 2.02 |
| *P. teleius* | PL-T03 | 27.07.2020 | 5.261042 | 46.005833 | PontLoup | 960530 | 8016 | 3.58 |
| *P. teleius* | PL-T04 | 27.07.2020 | 5.261066 | 46.005894 | PontLoup | 1601830 | 8119 | 2.35 |
| *P. teleius* | PL-T05 | 27.07.2020 | 5.260981 | 46.005929 | PontLoup | 1807396 | 8173 | 1.70 |
| *P. teleius* | PL-T06 | 27.07.2020 | 5.260849 | 46.006103 | PontLoup | 944836 | 7875 | 5.28 |
| *P. teleius* | PL-T07 | 27.07.2020 | 5.26129 | 46.00615 | PontLoup | 1542808 | 8236 | 0.94 |
| *P. teleius* | PL-T08 | 27.07.2020 | 5.261361 | 46.006021 | PontLoup | 1231172 | 8038 | 3.32 |
| *P. teleius* | PL-T09 | 27.07.2020 | 5.261699 | 46.005916 | PontLoup | 1583240 | 8187 | 1.53 |
| *P. teleius* | PL-T10 | 27.07.2020 | 5.261936 | 46.006117 | PontLoup | 585602 | 6487 | 21.98 |
| *P. teleius* | PL-T11 | 27.07.2020 | 5.262962 | 46.005649 | PontLoup | 1188148 | 8082 | 2.79 |
| *P. teleius* | PL-T12 | 27.07.2020 | 5.263916 | 46.005831 | PontLoup | 1626382 | 8085 | 2.75 |
| *P. teleius* | Vx-T01 | 11.08.2020 | 5.59453 | 45.954947 | VauxMelogne | 863850 | 6668 | 19.80 |
| *P. teleius* | Vx-T02 | 11.08.2020 | 5.590637 | 45.953046 | VauxMelogne | 1782990 | 8182 | 1.59 |
| *P. teleius* | Vx-T03 | 11.08.2020 | 5.590744 | 45.952979 | VauxMelogne | 1728980 | 8076 | 2.86 |

**Supplementary Table 4.** Pairwise differentiation level $F_{ST}$ and migration rates estimated between each locality of the three species, including the Jost's D, Nei's Gst, and the effective number of migrants *N*m. Values in bold indicate reliable estimates with sample sizes greater than 10.

| Species | loc1 | loc2 | n loc1 | n loc2 | FST | 1>2 | 1<2 | 1>2 | 1<2 | 1>2 | 1<2 |
|---|---|---|---|---|---|---|---|---|---|---|---|
| | Belloire | Bidonnes | 12 | 4 | 0.614 | 0.125 | 0.145 | 0.380 | 0.523 | 0.377 | 0.519 |
| | **Belloire** | **Cerin** | **12** | **12** | **0.536** | **0.376** | **0.369** | **0.609** | **0.522** | **0.607** | **0.521** |
| *P. alcon* | **Belloire** | **Ormes** | **12** | **12** | **0.581** | **0.235** | **1.000** | **0.373** | **1.000** | **0.371** | **1.000** |
| | Bidonnes | Cerin | 4 | 12 | 0.627 | 0.144 | 0.127 | 0.527 | 0.364 | 0.523 | 0.361 |
| | Bidonnes | Ormes | 4 | 12 | 0.752 | 0.069 | 0.119 | 0.392 | 0.536 | 0.385 | 0.530 |
| | **Cerin** | **Ormes** | **12** | **12** | **0.646** | **0.160** | **0.444** | **0.329** | **0.827** | **0.326** | **0.825** |
| | **Bidonnes** | **Broues** | **12** | **12** | **0.157** | **0.320** | **0.256** | **1.000** | **0.943** | **1.000** | **0.943** |
| | **Bidonnes** | **Epierre** | **12** | **11** | **0.506** | **0.022** | **0.089** | **0.120** | **0.231** | **0.119** | **0.231** |
| | Bidonnes | Flon | 12 | 2 | 0.168 | 0.112 | 0.254 | 0.330 | 0.615 | 0.330 | 0.615 |
| | **Bidonnes** | **Intriat** | **12** | **12** | **0.533** | **0.022** | **0.096** | **0.113** | **0.189** | **0.112** | **0.189** |
| *P. nausithous* | **Broues** | **Epierre** | **12** | **11** | **0.492** | **0.021** | **0.107** | **0.116** | **0.231** | **0.115** | **0.231** |
| | Broues | Flon | 12 | 2 | 0.137 | 0.117 | 0.284 | 0.366 | 0.677 | 0.365 | 0.677 |
| | **Broues** | **Intriat** | **12** | **12** | **0.519** | **0.020** | **0.113** | **0.113** | **0.209** | **0.112** | **0.209** |
| | Epierre | Flon | 11 | 2 | 0.568 | 0.036 | 0.020 | 0.145 | 0.124 | 0.144 | 0.123 |
| | **Epierre** | **Intriat** | **11** | **12** | **0.226** | **0.604** | **1.000** | **0.574** | **0.616** | **0.574** | **0.617** |
| | Flon | Intriat | 2 | 12 | 0.610 | 0.020 | 0.040 | 0.124 | 0.135 | 0.123 | 0.134 |
| | **Bidonnes** | **Broues** | **20** | **12** | **0.093** | **0.427** | **0.492** | **0.933** | **1.000** | **0.933** | **1.000** |
| | **Bidonnes** | **Epierre** | **20** | **13** | **0.494** | **0.024** | **0.124** | **0.071** | **0.116** | **0.071** | **0.116** |
| | Bidonnes | Flon | 20 | 4 | 0.145 | 0.186 | 0.552 | 0.284 | 0.553 | 0.283 | 0.553 |
| | Bidonnes | Lavours | 20 | 5 | 0.276 | 0.101 | 0.159 | 0.191 | 0.204 | 0.191 | 0.203 |
| | Bidonnes | Melogne | 20 | 4 | 0.377 | 0.029 | 0.249 | 0.074 | 0.190 | 0.073 | 0.190 |
| | **Bidonnes** | **Pont Loup** | **20** | **12** | **0.467** | **0.029** | **0.122** | **0.082** | **0.125** | **0.081** | **0.125** |
| | Bidonnes | Vaux | 20 | 3 | 0.328 | 0.045 | 0.157 | 0.103 | 0.170 | 0.103 | 0.170 |
| | **Broues** | **Epierre** | **12** | **13** | **0.537** | **0.023** | **0.112** | **0.064** | **0.108** | **0.064** | **0.108** |
| | Broues | Flon | 12 | 4 | 0.155 | 0.188 | 0.436 | 0.268 | 0.489 | 0.268 | 0.489 |
| | Broues | Lavours | 12 | 5 | 0.288 | 0.105 | 0.151 | 0.175 | 0.186 | 0.175 | 0.186 |
| | Broues | Melogne | 12 | 4 | 0.407 | 0.028 | 0.211 | 0.068 | 0.164 | 0.067 | 0.164 |
| | **Broues** | **Pont Loup** | **12** | **12** | **0.504** | **0.028** | **0.110** | **0.076** | **0.118** | **0.076** | **0.118** |
| | Broues | Vaux | 12 | 3 | 0.348 | 0.044 | 0.146 | 0.092 | 0.156 | 0.092 | 0.156 |
| *P. teleius* | Epierre | Flon | 13 | 4 | 0.646 | 0.043 | 0.021 | 0.072 | 0.060 | 0.071 | 0.059 |
| | Epierre | Lavours | 13 | 5 | 0.463 | 0.411 | 0.055 | 0.165 | 0.080 | 0.165 | 0.079 |
| | Epierre | Melogne | 13 | 4 | 0.601 | 0.055 | 0.060 | 0.065 | 0.078 | 0.064 | 0.078 |
| | **Epierre** | **Pont Loup** | **13** | **12** | **0.374** | **0.546** | **0.303** | **0.151** | **0.131** | **0.151** | **0.131** |
| | Epierre | Vaux | 13 | 3 | 0.549 | 0.100 | 0.051 | 0.093 | 0.081 | 0.093 | 0.081 |
| | Flon | Lavours | 4 | 5 | 0.363 | 0.110 | 0.071 | 0.148 | 0.110 | 0.148 | 0.110 |
| | Flon | Melogne | 4 | 4 | 0.533 | 0.026 | 0.069 | 0.063 | 0.105 | 0.063 | 0.105 |
| | Flon | Pont Loup | 4 | 12 | 0.612 | 0.026 | 0.045 | 0.066 | 0.074 | 0.066 | 0.074 |
| | Flon | Vaux | 4 | 3 | 0.440 | 0.044 | 0.058 | 0.089 | 0.099 | 0.088 | 0.099 |
| | Lavours | Melogne | 5 | 4 | 0.278 | 0.087 | 1.000 | 0.106 | 0.347 | 0.106 | 0.348 |
| | Lavours | Pont Loup | 5 | 12 | 0.395 | 0.082 | 0.374 | 0.106 | 0.179 | 0.106 | 0.179 |
| | Lavours | Vaux | 5 | 3 | 0.144 | 0.202 | 0.585 | 0.201 | 0.415 | 0.201 | 0.415 |
| | Melogne | Pont Loup | 4 | 12 | 0.551 | 0.087 | 0.058 | 0.086 | 0.062 | 0.086 | 0.062 |
| | Melogne | Vaux | 4 | 3 | 0.207 | 0.896 | 0.184 | 0.377 | 0.154 | 0.377 | 0.154 |
| | Pont Loup | Vaux | 12 | 3 | 0.487 | 0.110 | 0.075 | 0.091 | 0.091 | 0.091 | 0.091 |
